# Clearing spheroids for 3D fluorescent microscopy: combining safe and soft chemicals with deep convolutional neural network

**DOI:** 10.1101/2021.01.31.428996

**Authors:** Ali Ahmad, Saba Goodarzi, Carole Frindel, Gaëlle Recher, Charlotte Riviere, David Rousseau

## Abstract

In life sciences, there are increasing interest in 3D culture models to better reproduce the 3D environment encountered in-vivo. Imaging of such 3D culture models is instrumental for drug discovery, but face several issues before its use becomes widespread. Extensive microscopic investigation of these 3D cell models faces the challenge of light penetration in depth in opaque biological tissues. To overcome this limit, diverse clearing techniques have emerged over the past decades. However, it is not straightforward to choose the best clearing protocols, and assess quantitatively their clearing efficiency. Focusing on spheroids, we propose a combination of fast and cost-effective clearing procedure for such medium-sized samples. A generic method with local contrast metrics and deep convolutional neural network-based segmentation of nuclei is proposed to quantify the efficiency of clearing. We challenged this method by testing the possibility to transfer segmentation knowledge from a clearing protocol to another. The later results support the pertinence of training deep learning algorithms on cleared samples to further use the segmentation pipeline on non-cleared ones. This second step of the protocol gives access to digital clearing possibilities applicable to live and high-throughput optical imaging.

## Introduction

Spheroids are three-dimensional (3D), heterogeneous aggregates of proliferating, quiescent and necrotic in vitro cell culture systems [1]. They have gained increasing interest in drug screening because of their ability to closely mimic the main features of physiological cell to cell and cell to matrix contacts as in in vivo human solid tumors [2, 3]. Because of the standardization of their production, spheroids became a model of choice in the context of the 3R (replace, reduce, refine) notably for high-throughput drug screening [4, 5]. Yet, the experimental readouts are often quantitative, between each size-controlled spheroid, but fail at providing insights in regards with single-cell information within the spheroid itself, due to its lack of amenability for in-depth microscopy. One main challenge is the visualization of spheroids via fluorescence imaging due to the light scattering inducing limited depth penetration in such opaque structure and therefore low data analysis performance at a single-cell level [6, 7]. One solution to enhance visualization of fluorescently-labelled spheroids is to use optical clearing techniques, which improve in-depth imaging and allow high quality image acquisition and high-throughput image analysis. The rough principle of optical clearing is to uniformise the optical index within the 3D structure, so as to minimise diffraction. One of the option is to remove the lipids that contribute to light scattering effects, or, to replace water by a solution closer to the lipid refractive index [8–13]. So far, assessment of the quality of clearing in 3D spheroids has been performed with local quality metrics such as signal to noise ratio (SNR) and contrast to noise ratio (CNR) [14–16]. However, it is not straight-forward to relate local metrics with final quantitative measures of interest. We not only propose to revisit the assessment of clearing quality in such a perspective, but we also provide an assessment based on segmentation metrics.

Final quantitative measures for high-throughput image analysis of spheroids mostly rely on whole-spheroid fluorescent measures, size analysis and matrix invasion as an indicator of drug anti metastatic effects [17–22]. These metrics provide useful macroscopic quantitative information but do not provide enough biological details at single cell level for assessing therapy responses nor quantitative analysis such as cell counting and aggregation studies, and also no characteristics over time such as growth ratio and proliferation ratio. To overcome these limitations, individual cell nuclei segmentation methods have been developed for 2D and 3D spheroid images [23–30]. State-of-the-art methods for cell nuclei segmentation in microscopy are currently the deep learning-based methods. Also, a specific interest of the deep learning-based approach, by contrast with a standard image processing pipeline composed of denoising step [31] followed by a segmentation step [32, 33], is to offer an end-to-end learning process where all the pipeline is optimized at the same time. Several architectures were developed to segment nuclei by integrating two or more channels in the output of deep learning architecture or by applying post processing methods to the predicted segmentation maps to enhance segmentation quality [34–40]. Deep learning methods were applied to segment entire spheroids of different sizes, shapes, and illumination conditions [41] and also to segment nuclei of 3D spheroid images [42]. In these closely related work, only one clearing method was investigated. By contrast in this article we propose, with a deep learning perspective, a protocol to compare clearing methods for the segmentation of nuclei in spheroids under 3D fluorescence microscopy.

A scheme of the operating pipeline of the article is provided in Figure 1. We compare clearing protocols RapiClear and Glycerol (Figure 1.A) to non cleared samples (Control). The two clearing methods investigated have been chosen for their simplicity and non-toxicity. Both Rapiclear [43] and Glycerol [15] have already been reported as fast and cost-effective yet efficient clearing procedures, for medium-sized samples such as organoids and spheroids of few hundreds of micrometers. First, we assess the clearing quality of spheroid images acquired with confocal fluorescence microscopy with the conventional local metrics such as signal to noise ratio (SNR) and contrast to noise ratio (CNR) (Figure 1.B). Then, we use deep learning segmentation methods to assess the quality of the clearing (Figure 1.C,D). Finally, we investigate the possibility of digital clearing of non-cleared data with cleared data models via transfer learning and the transferability of knowledge from a clearing protocol to another (Figure 1.E). In this manuscript, we show that using a simple deep learning strategy, it is possible to get reliable segmented images even for images with low intensity and low signal-to-noise ratio. Also, we demonstrate the segmentation knowledge transferability from cleared samples to native tissues for fast digital-clearing of living specimens on the fly.

**Fig 1.**
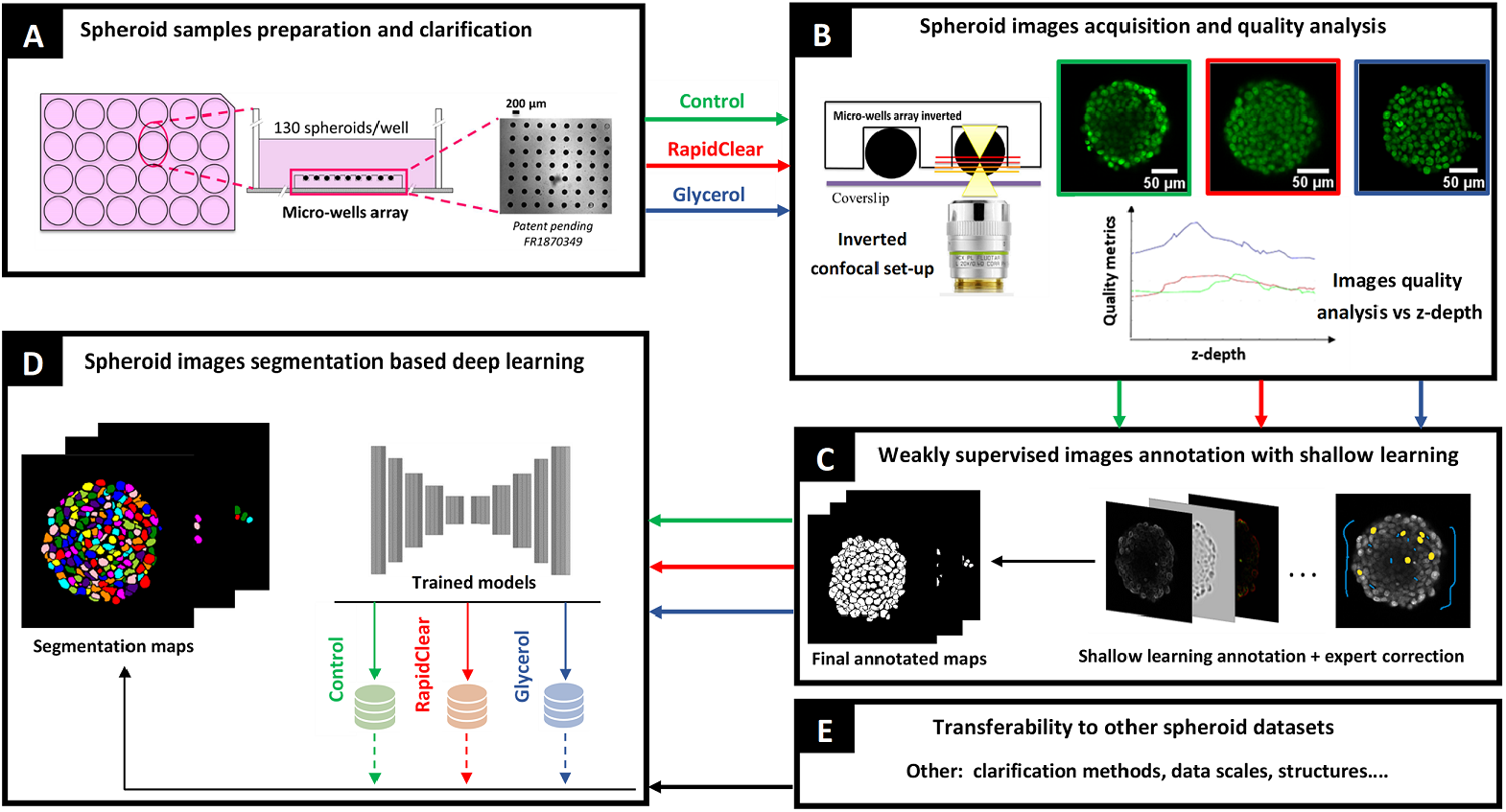
Protocol pipeline: **(A)** Spheroid samples are cultured in micro-well arrays. Control are the non cleared samples and others are clarified with two distinct clearing methods Rapiclear and Glycerol. **(B)** Then, spheroid images acquired with a confocal microscope are analyzed for quality assessment along z-depth. **(C)** A dataset of images for each clarification method is annotated using shallow learning and corrected by an expert. **(D)** Finally, deep learning models are trained with the annotated data sets and the three segmentation models trained on Rapiclear, Control and Glycerol are tested for **(E)** data transferability on various clarification methods.

## Results and Discussion

### Comparison of clearing methods with local metrics

We assessed the efficiency of RapiClear and Glycerol clearing protocols on image quality by comparing their datasets with non cleared datasets (Control) using image local quality metrics evaluation. This was quantitatively evaluated in depth on the 3D spheroid stacks (z-depth) based on the computation of various local metrics using patches cropped from the center of the spheroid signal in each slice (see SI Figures 1.A). The used metrics were the signal to noise ratio (SNR) (Equation 1) and the contrast to noise ratio (CNR) calculated by two ways: from Bhattacharyya coefficient (BC) (Equation 2) and from Fisher ratio (FR) (Equation 3). Figure 2 shows the xz planes at the center of spheroids and the xy planes at selected depth (from 70% to 100% of maximum diameter, corresponding to the range of depth where the three conditions can be compared) also the mean and the standard deviation (std) of the normalized average intensity (average I/Imax) evolution and of the computed local metrics from three spheroids for each Control, RapiClear and Glycerol conditions. Visual qualitative inspection of the xz and xy planes of spheroids are provided in Fig 2.A,B. Quantitative evaluation of image quality (Fig 2.C) shows the degradation of image with depth z.

**Fig 2.**
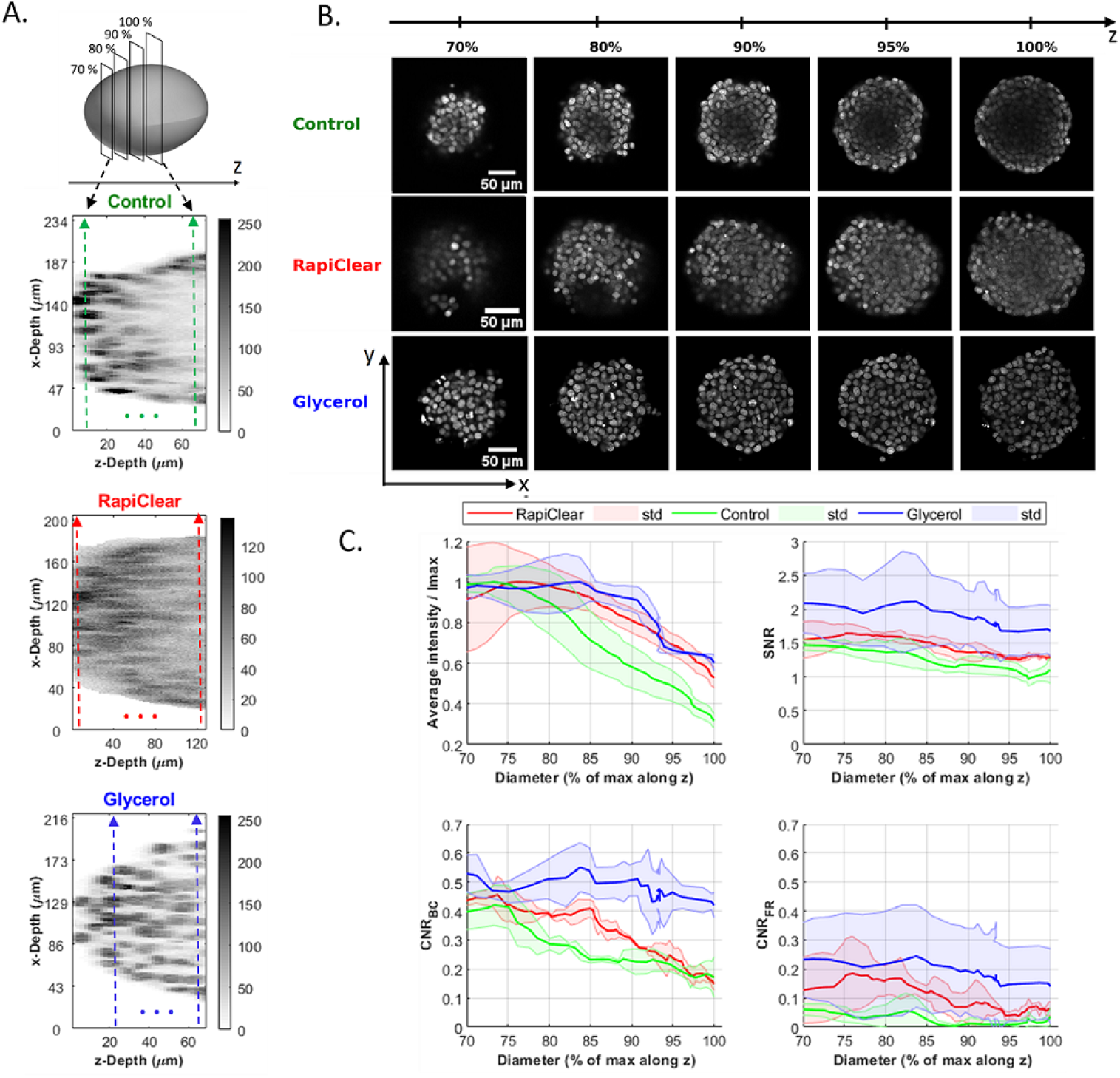
Image quality analysis function of the depth (*z*) defined as the percentage of the maximum diameter for the datasets produced with Control, RapiClear and Glycerol clarification methods. Slices with 100% of diameter are considered as the center of the 3D spheroids. **(A)** Illustration of the *xz* slices. Colored dashed lines correspond to the location (in *μm*) of the slices *xy* with 70% and 100% of maximum diameter. **(B)** Visualization of test images (xy slices) normalized in depths (*z*) and used to evaluate segmentation performance for each clarification method. **(C)** The mean and standard deviation (std) of the average intensity, SNR, *CNR_BC_* by Battacharayya and *CNR_FR_* by Fisher ratio computed for 3 spheroids for each clarification method and plotted as a function of the percentage of maximum diameter.

Concerning intensities, an important drop is recorded for all clearing methods. This drop is more pronounced in the non cleared (control) spheroids. The intensity evolutions with depth are similar for Rapiclear and Glycerol. SNR and FR metrics appear almost constant for the three investigated clearing conditions while BC shows global degradation along depth z. For the three metrics SNR, FR and BC, Glycerol shows significantly better values by comparison with Rapiclear and Control. However, it is uneasy to produce a secured prediction of the effect of clearing along z with local metrics. This is why it was important also to confront what would be found with conventional local metrics with a machine learning perspective.

### Spheroid 2D images segmentation

We used deep learning segmentation methods to characterize the quality of spheroid clearing protocols. Several segmentation methods such as Unet, Dist and Stardist, associated with post-processing steps based on dynamic morphology reconstruction and watershed algorithm (DM) were applied to the test images of Figure 2.B produced from the tested clearing conditions. We used the F1-score to quantify the quality of segmentation as a pixel wise metric and the aggregated Jaccard Index (AJI) as an object wise metric. We tested the segmentation efficiency for two cases: (i) when test images and pre-trained segmentation models are from the same clearing method and (ii) in a cross way taking test images from a clearing condition and using pre-trained model from another. Table 1 shows the mean values and the standard deviations (std) of the F1-score and AJI computed for test images located at 70, 80, 90, 95 and 100% of maximum diameter along z.

**Table 1.** Mean and standard deviation of the computed segmentation metric (F1-score, AJI) for five test images produced from Control, RapiClear and Glycerol. Tests are realized in cross way between the segmentation models (A,B and C). And also for five test images produced from TDE, ScaleS and CUBIC to test data transferability from a clearing protocol to another clearing protocol (D, E and F). Various segmentation methods are used. The bold values are best segmentation model for each clearing method datasets. Except for Cubic clearing method where all methods provide low performance, Glycerol appears as the best method on which one should train the deep learning models in order to benefit from digital clearing.

| Test data | Segmentation models | F1-score ± std | | | AJI ± std | | | | |
| --- | --- | --- | --- | --- | --- | --- | --- | --- | --- |
|  |  | *Unet* | *Dist* | *Stardist* | *Unet* | *Unet+DM* | *Dist* | *Dist + DM* | *Stardist* |
| Control<br>(A) | Control<br>(1) | 0.85<br>$\pm 0.01$ | 0.85<br>$\pm 0.01$ | 0.84<br>$\pm 0.01$ | 0.07<br>$\pm 0.06$ | 0.51<br>$\pm 0.03$ | 0.26<br>$\pm 0.05$ | 0.52<br>$\pm 0.03$ | 0.59<br>$\pm 0.03$ |
| | RapiClear<br>(2) | 0.84<br>$\pm 0.02$ | 0.85<br>$\pm 0.01$ | 0.72<br>$\pm 0.04$ | 0.14<br>$\pm 0.03$ | 0.52<br>$\pm 0.05$ | 0.16<br>$\pm 0.03$ | 0.52<br>$\pm 0.03$ | 0.42<br>$\pm 0.07$ |
|  | **Glycerol**<br>(3) | **0.86**<br>**$\pm 0.02$** | **0.86**<br>**$\pm 0.01$** | **0.84**<br>**$\pm 0.02$** | **0.01**<br>**$\pm 0.01$** | **0.57**<br>**$\pm 0.05$** | **0.06**<br>**$\pm 0.04$** | **0.55**<br>**$\pm 0.02$** | **0.55**<br>**$\pm 0.06$** |
| RapiClear<br>(B) | Control<br>(1) | 0.82<br>$\pm 0.01$ | 0.83<br>$\pm 0.01$ | 0.81<br>$\pm 0.01$ | 0.03<br>$\pm 0.02$ | 0.41<br>$\pm 0.02$ | 0.07<br>$\pm 0.03$ | 0.47<br>$\pm 0.05$ | 0.52<br>$\pm 0.03$ |
| | RapiClear<br>(2) | 0.83<br>$\pm 0.01$ | 0.84<br>$\pm 0.01$ | 0.76<br>$\pm 0.03$ | 0.11<br>$\pm 0.09$ | 0.46<br>$\pm 0.04$ | 0.08<br>$\pm 0.04$ | 0.48<br>$\pm 0.02$ | 0.43<br>$\pm 0.06$ |
|  | **Glycerol**<br>(3) | **0.83**<br>**$\pm 0.03$** | **0.83**<br>**$\pm 0.02$** | **0.80**<br>**$\pm 0.03$** | **0.01**<br>**$\pm 0.01$** | **0.46**<br>**$\pm 0.02$** | **0.03**<br>**$\pm 0.01$** | **0.47**<br>**$\pm 0.03$** | **0.49**<br>**$\pm 0.03$** |
| Glycerol<br>(C) | Control<br>(1) | 0.89<br>$\pm 0.01$ | 0.89<br>$\pm 0.01$ | 0.89<br>$\pm 0.01$ | 0.29<br>$\pm 0.01$ | 0.63<br>$\pm 0.01$ | 0.46<br>$\pm 0.06$ | 0.64<br>$\pm 0.03$ | 0.68<br>$\pm 0.02$ |
| | RapiClear<br>(2) | 0.88<br>$\pm 0.01$ | 0.87<br>$\pm 0.01$ | 0.78<br>$\pm 0.02$ | 0.40<br>$\pm 0.07$ | 0.63<br>$\pm 0.03$ | 0.46<br>$\pm 0.04$ | 0.62<br>$\pm 0.02$ | 0.51<br>$\pm 0.03$ |
|  | **Glycerol**<br>(3) | **0.91**<br>**$\pm 0.01$** | **0.91**<br>**$\pm 0.01$** | **0.90**<br>**$\pm 0.01$** | **0.34**<br>**$\pm 0.07$** | **0.70**<br>**$\pm 0.02$** | **0.47**<br>**$\pm 0.05$** | **0.71**<br>**$\pm 0.02$** | **0.70**<br>**$\pm 0.02$** |
| TDE<br>(D) | Control<br>(1) | 0.88<br>$\pm 0.01$ | 0.88<br>$\pm 0.02$ | 0.84<br>$\pm 0.02$ | 0.02<br>$\pm 0.01$ | 0.44<br>$\pm 0.07$ | 0.06<br>$\pm 0.05$ | 0.56<br>$\pm 0.05$ | 0.59<br>$\pm 0.04$ |
| | RapiClear<br>(2) | 0.89<br>$\pm 0.01$ | 0.89<br>$\pm 0.02$ | 0.78<br>$\pm 0.02$ | 0.08<br>$\pm 0.04$ | 0.60<br>$\pm 0.01$ | 0.09<br>$\pm 0.04$ | 0.60<br>$\pm 0.02$ | 0.52<br>$\pm 0.06$ |
|  | **Glycerol**<br>(3) | **0.90**<br>**$\pm 0.02$** | **0.89**<br>**$\pm 0.03$** | **0.84**<br>**$\pm 0.02$** | **0.02**<br>**$\pm 0.02$** | **0.39**<br>**$\pm 0.05$** | **0.02**<br>**$\pm 0.02$** | **0.54**<br>**$\pm 0.02$** | **0.58**<br>**$\pm 0.04$** |
| ScaleS<br>(E) | Control<br>(1) | 0.85<br>$\pm 0.03$ | 0.86<br>$\pm 0.03$ | 0.83<br>$\pm 0.03$ | 0.29<br>$\pm 0.13$ | 0.54<br>$\pm 0.03$ | 0.39<br>$\pm 0.1$ | 0.59<br>$\pm 0.05$ | 0.62<br>$\pm 0.05$ |
| | RapiClear<br>(2) | 0.83<br>$\pm 0.06$ | 0.85<br>$\pm 0.05$ | 0.77<br>$\pm 0.06$ | 0.22<br>$\pm 0.1$ | 0.58<br>$\pm 0.03$ | 0.26<br>$\pm 0.13$ | 0.58<br>$\pm 0.06$ | 0.50<br>$\pm 0.06$ |
|  | **Glycerol**<br>(3) | **0.87**<br>**$\pm 0.03$** | **0.85**<br>**$\pm 0.03$** | **0.85**<br>**$\pm 0.03$** | **0.12**<br>**$\pm 0.07$** | **0.57**<br>**$\pm 0.04$** | **0.14**<br>**$\pm 0.11$** | **0.56**<br>**$\pm 0.05$** | **0.63**<br>**$\pm 0.03$** |
| CUBIC<br>(F) | **Control**<br>(1) | **0.68**<br>**$\pm 0.002$** | **0.69**<br>**$\pm 0.03$** | **0.24**<br>**$\pm 0.04$** | **0.003**<br>**$\pm 0.001$** | **0.33**<br>**$\pm 0.04$** | **0.004**<br>**$\pm 0.001$** | **0.16**<br>**$\pm 0.04$** | **0.07**<br>**$\pm 0.02$** |
| | RapiClear<br>(2) | 0.66<br>$\pm 0.01$ | 0.63<br>$\pm 0.02$ | 0.42<br>$\pm 0.02$ | 0.003<br>$\pm 0.001$ | 0.23<br>$\pm 0.03$ | 0.003<br>$\pm 0.001$ | 0.19<br>$\pm 0.02$ | 0.11<br>$\pm 0.001$ |
| | Glycerol<br>(3) | 0.58<br>$\pm 0.01$ | 0.58<br>$\pm 0.01$ | 0.26<br>$\pm 0.04$ | 0.002<br>$\pm 0.001$ | 0.23<br>$\pm 0.04$ | 0.002<br>$\pm 0.001$ | 0.16<br>$\pm 0.01$ | 0.06<br>$\pm 0.01$ |

Interestingly, in accordance with the observation recorded by the local quality metrics, Glycerol clearing protocol provides the best segmentation performance in all tested configurations (Table 1.C-3). In addition, it is remarkable that Glycerol segmentation model is also powerful to segment data sets from Control or from Rapiclear clearing when segmentation method is properly chosen (Table 1.A-3,B-3). It is also striking to note that segmentation results of Control and RapiClear test images gives higher F1-score and AJI values with segmentation models trained on Glycerol than with models trained on their data. As shown in Fig. 3.A, these average results are almost stationary along z for all clearing methods tested (also see qualitative illustration of the final segmentation in Figure 3.B and SI Figures 2 and 3.A). This demonstrates that the local evolution in intensity or SNR shown in Figure 2 do not necessarily correlate with the segmentation performances.

**Fig 3.**
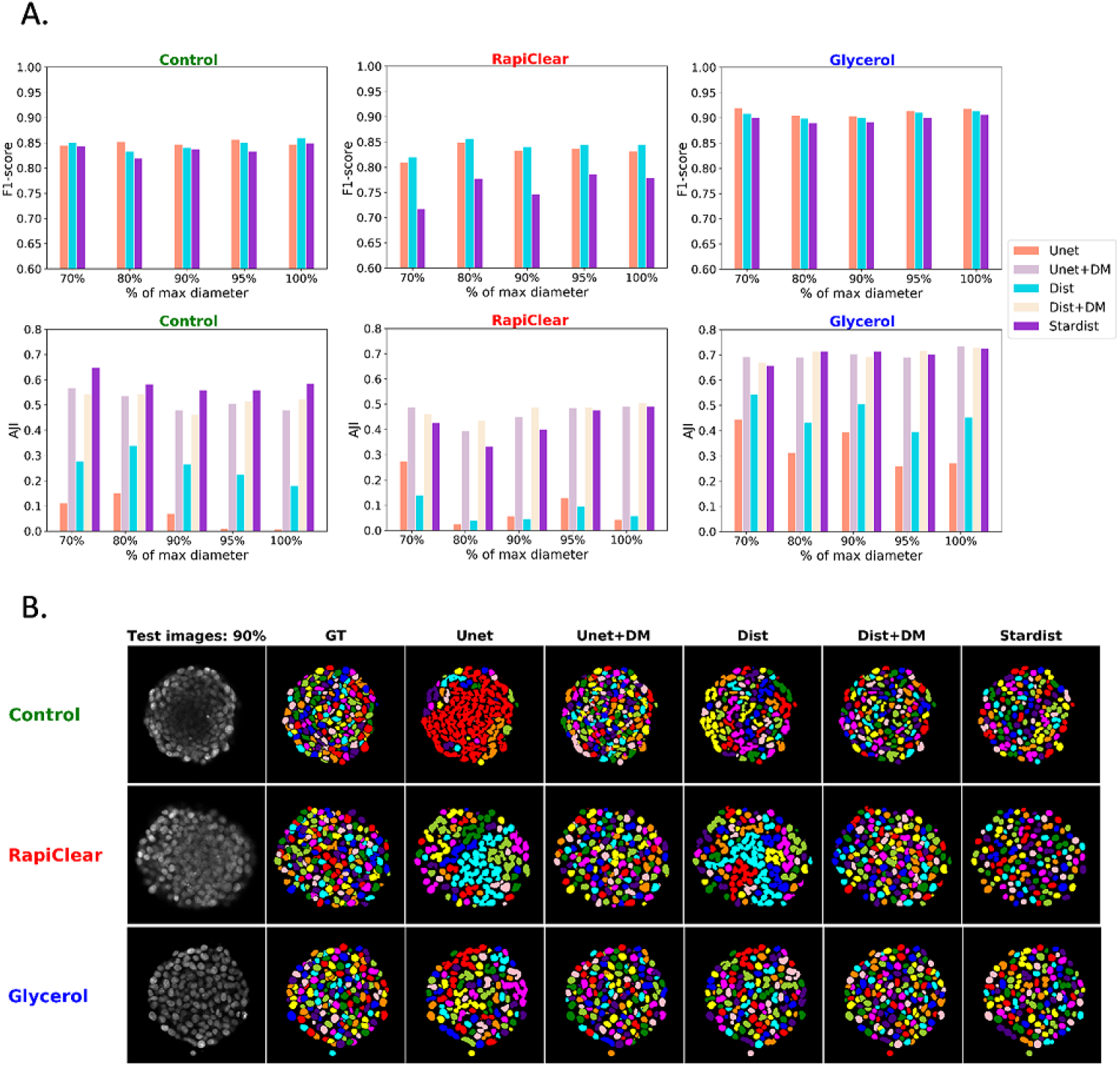
Segmentation results as a function of depths (*z*). **(A)** Quantitative segmentation results (F1-score, AJI) for the five test images taken at depth ranging from 70, 80, 90, 95 and 100% defined as the percentage of maximum diameter. Segmentation methods are trained and tested on the same clearing method. **(B)** Qualitative illustration of the final segmentation of the slices at 90% of maximum diameter produced from Control, RapiClear and Glycerol clearing methods.

### Segmentation models transferability to other data sets

Finally, we tested the transferability from cleared datasets to another datasets by segmenting test images from another clearing methods, another structure and also another imaging system using pre-trained segmentation models of Control, Rapiclear and Glycerol. For this study, we used colorectal carcinoma HCT-116 cell lines test images cleared with TDE and acquired with a laser scan confocal system (Nikon). Also, we selected two datasets from the literature such as Breast carcinoma T47D cells cleared with ScaleS method and acquired with a spinnig disk confocal system (Opera Phenix) [23] and colorectal carcinoma HCT-116 cell line cleared with CUBIC method and acquired with a light-sheet fluorescence (LSF) microscope [44]. Table 1.D,E,F shows the mean and standard deviation of computed F1-score and AJI segmentation metrics for each test images produced from TDE, ScaleS and CUBIC clearing methods and segmented with the Control, RapiClear and Glycerol pre-trained models. The segmentation results are high for both TDE and ScaleS (see Table 1 and Figure 4.A). Similarly to what was found for Rapiclear and Control, Glycerol appears, for both TDE and ScaleS also, as the best method to train on to benefit from transfer learning. This experiment clearly proves the feasibility of information transferability by digital clearing from clearing method to another acquired with the same imaging system but with different structure and data scale. Contrariwise, CUBIC dataset acquired with light-sheet fluorescence (LSF) microscopy and presenting very different contrast and artefacts shows low performance. This experiment points the limit of data transferability by digital clearing (see Figure 4.B and SI Figures 3.B and 4 for qualitative illustration of final segmentation maps). Figure 5.A illustrates the qualitative interest of digital clearing available at the output of neural networks when the best segmentation neural network (Dist segmentation) is trained on the best clearing method (Glycerol) and applied to uncleared data or cleared with Rapiclear. It is obvious with Fig. 5.A that the intensity attenuation recorded with local metrics along z can be compensated with the deep neural network thanks to digital clearing. As shown in Fig. 5.B in comparison with ground truth, a significant improvement of the counting of cells along z is brought by training on images obtained with the best clearing method while applying the model directly on control or on a less efficient clearing method. This opens also the possibility of an efficient post-segmentation 3D reconstruction of full spheroids (see SI Figure 5 and SI videos).

**Fig 4.**
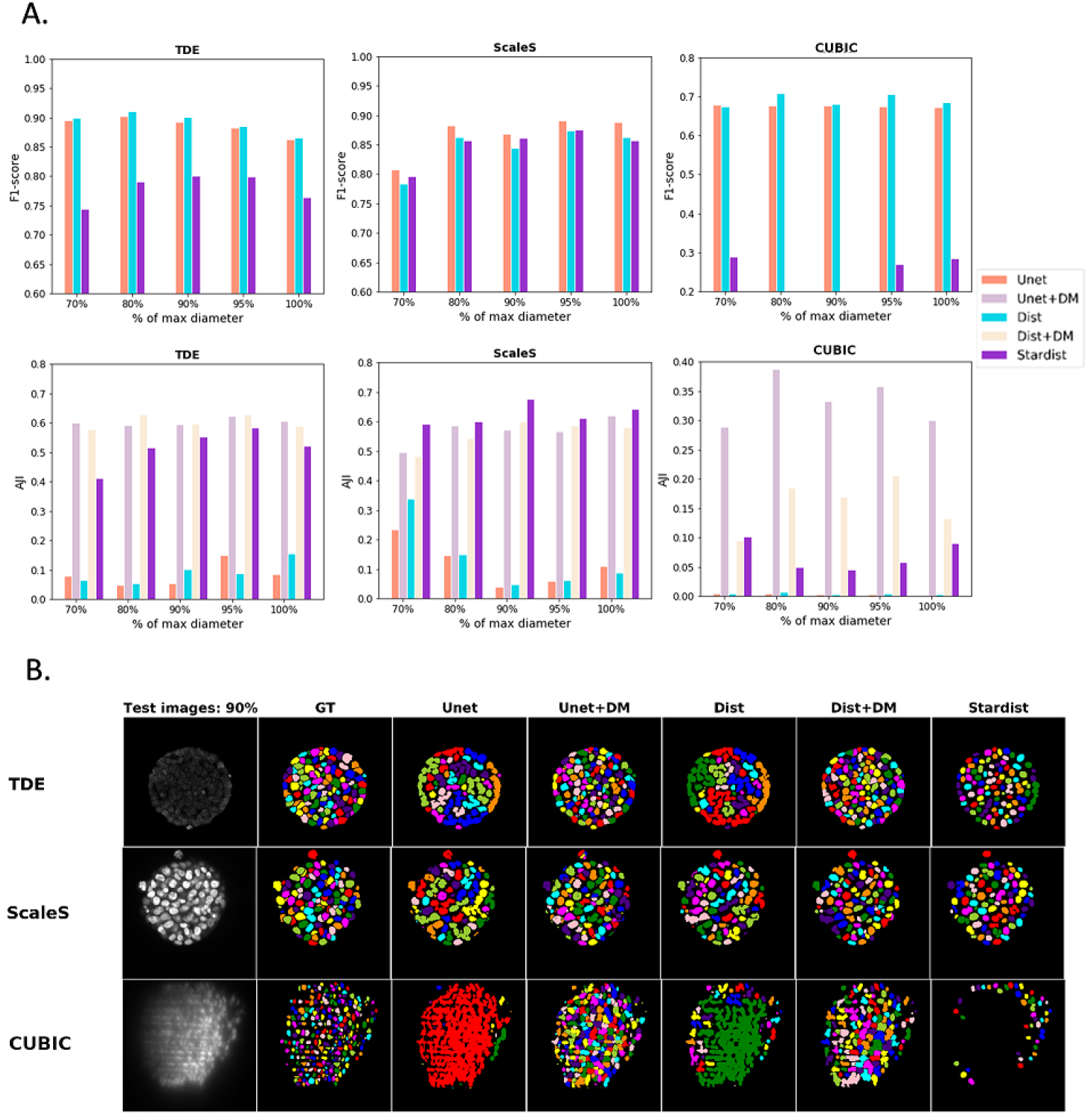
Data transferability assessment: segmentation results as a function of depths (z) defined as the percentage of maximum diameter. **(A)** Quantitative segmentation results (F1-score, AJI) for the five test images produced from TDE, ScaleS and CUBIC clearing methods and segmented using the RapiClear, Glycerol and Control segmentation models respectively. **(B)** Qualitative illustration of the final segmentation results for the slices at 90% of maximum diameter.

**Fig 5.**
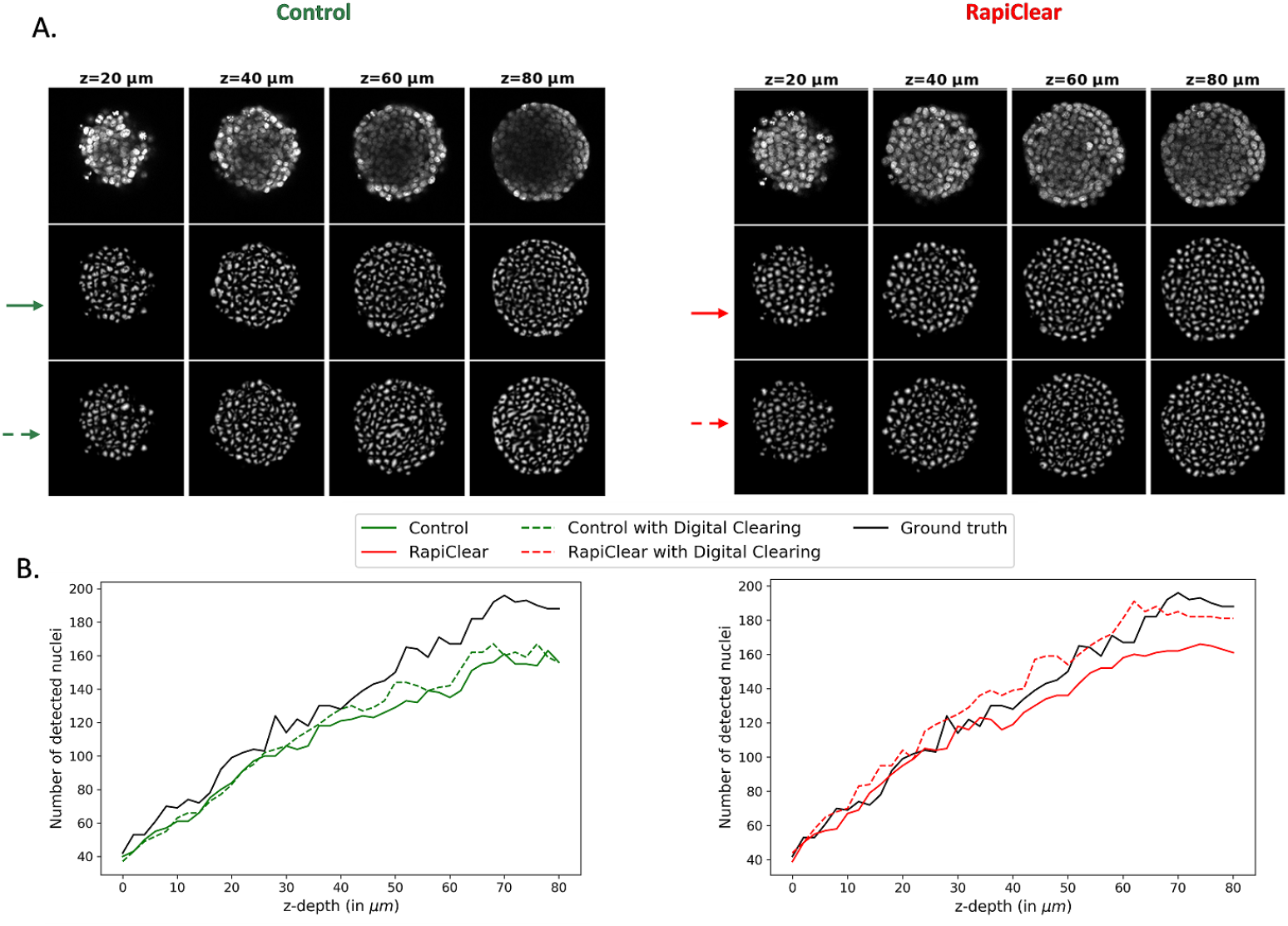
Digital clearing effect analysis. **(A)** Slices of non cleared spheroid (control) and same spheroid cleared with RapiClear are illustrated function of z-depth (*μm*). The predicted distance maps at the output of Unet architecture for Dist segmentation method computed for the slices with control model (green solid line) and RapiClear model (red solid line) and also with digital clearing via glycerol model (green and red dashed lines). **(B)** Number of detected nuclei computed after Dist segmentation function of the depth z in *μm* for the same non cleared and cleared spheroid before and after digital clearing. Ground truth in black solid line corresponds to a manual counting by an expert.

## Conclusion

In this report, we have investigated the interest of a deep learning perspective for the comparison of clearing methods in fluorescence microscopy. This was illustrated for a task of nuclei segmentation in spheroids under 3D fluorescent microscopy. The best clearing method identified with conventional local metric follows the performance of state-of-the-art deep learning based segmentation methods. However, because the training datasets include images from various depth, no real influence of depth was observed on segmentation performance contrarily to what was found with local metric. This demonstrates that local metric can be used to select an optimal clearing method but that final performances can be made invariant to the remaining noise thanks to adequate machine learning strategies.

We specially tested the interest of transfer learning in this context. We investigated the interest to train on one clearing condition and to test on another. The best segmentation method was also found to be the one which gave the best transfer results. This is specially interesting when the test images are produced in uncleared conditions. Training a model on the best cleared conditions enable to perform segmentation of nuclei in non cleared conditions with enhanced performances. This opens the way to digital clearing of the samples. Annotated cleared images of high quality produced with time consuming protocols could enable anyone to denoise images acquired with much simpler protocols. For this reason we release our annotated data set publicly as pointed in section S1 Dataset.

These results could be extended in various directions. The inference of the proposed segmentation approach is very fast (around 2 minutes for a stack of 41 slices of 40 Mbs with a processor Intel Core i7-6700HQ CPU @ 2.60 GHz). It can be easily used for on the fly segmentation during acquisition for fixed samples, as well as for live imaging (for experiments where samples are acquired over extended time-period [tens of hours], but with rather long time intervals [higher then 10 min]). By bringing such single cell metrics directly available to the user, it opens up promising applications to screen therapeutics drugs within 3D environment closer to in-vivo. Colorectal cancer spheroids were used here as an example. The method is also readily available for organoïds that have emerged as very powerful in vitro models to mimic various normal and pathological situations [45, 46]. The method can also be easily extended to the segmentation of other biological structures of interest, such as the overall cell shapes, as long as the manually segmented datasets is implemented.Other clarification methods specially optimized for thick samples [15,47–49] could then be also tested with the same global approach presented in this article. The most promising result lay in the possibility of digital clearing. In this study we performed it, for a first demonstration, by training directly on the best clearing method. Variants could be considered like style transfer [50, 51] or domain adaptation to further improve the digital clearing of the samples. Also, we demonstrated the possibility of digital clearing by transfer learning even when the two microscopes used are not strictly identical. This calls for a systematic quantitative study to assess the robustness of this finding with microscopes of various resolutions and aberrations.

## Materials and Methods

We present the details of all the steps described in the graphical abstract of Figure 1.

### Sample preparation and image acquisition

#### Cell lines and cell culture

Colorectal carcinoma cell line HCT-116 were used in this study. Cells were cultured in DMEM-GlutaMAX, supplemented with 10% of Heat-Inactivated Fetal Bovine Serum (FBS; Sigma, St. Louis, Missouri, US), 100 units / 100 *μg* of penicillin / streptomycin and passaged every 3-4 days.

#### Agarose-based micro-systems

The micro-systems consist of arrays of 130 micro-wells of 200 *μm* in diameter and height. They are used to create 130 reproducible spheroids per condition. This micro-system was produced by moulding agarose (Standard Agarose, 2% w/v) on PDMS molds created by photolithography process. A patent has been deposited for this process (FR 1870349).

#### Spheroid formation

Cells were seeded in each 24 wells-plate containing microsystems at a density of 1, 2.10^5^ cells/mL, 1 mL per well. They were placed under orbital agitation (160 rpm) for 2 hours in the incubator (37° C, 5% CO2) to increase cell sedimentation inside each micro-well. After 2 hours, the wells were rinsed with warm culture medium (3×) to get rid of cells that were out of the micro-wells. The 24 well-plate containing the microsystems were then kept in the incubator. Multicellular Tumor Spheroids (MCTs) are formed within 1 day and are used at Day 5 in this study.

#### Immunostaining

At day 5, cells were washed 3 times with warmed PBS for 5 min, followed by paraformaldehyde fixation (3.7% in PBS) for 20 minutes. All wells were then washed with PBS/3% BSA (3×5 min), permeabilized with 0.5% Triton for 20 min, and rinsed again with PBS/ 3% BSA (3x 5 min). To stain nucleus, NucGreen Dead 488 ReadyProbes Reagent were used (Invitrogen R37109, 2 drops/mL, i.e. 2 drops per wells, 4 hours at room temperature). The samples were washed with PBS (1×10 min) and kept protected from light in PBS at 4° C until image acquisition. Such non-cleared samples were either imaged directly (Control), or further processed with different clearing methods, described below.

#### Clarification

Different clarification techniques were used. For RapiClear-clarified samples, the microwells were incubated in RapiClear 1.52 solution (sunjinlab) overnight, then transferred in 0.5 mm Ispacer (sunjinlab, 2 spacers) and immerged in 35 *μL* of fresh RapiClear solution. Sealing was achieved using an additional sticky Ispacer and a coverslip. For Glycerol-clarified samples, the microwells were incubated in a 80% glycerol solution in PBS (v/v) overnight, then transferred in 0.5 mm Ispacer (sunjinlab, 2 spacers) and immerged in 35 *μL* of fresh 80% glycerol solution. The chemical compound 2,2’-thiodiethanol (TDE) was also tested in this study, following the procedure described in [52]. Briefly, the microwells were first incubated in a 20% TDEsolution in PBS (v/v) for 1 hour, then transferred in a 47% TDE solution in PBS (v/v) for 2 hours. The microwells were finally transferred in 0.5 mm Ispacer (sunjinlab, 2 spacers) and immerged in 35 *μL* of fresh 47% TDE/PBS solution. For ScaleS and CUBIC clarification methods, we used existing dataset of the literature described in [23, 44].

#### Image acquisition

A pile of z-stacks images of 1024 × 1024 pixels in xy was acquired using a classical confocal set-up. RapiClear images were acquired with Leica SP5 in resonant mode, a 20x dry objective lens (NA=0.7) is used with a pixel scale of 0.2 *μm* in xy and z-step 1 *μm*. Control, Glycerol and TDE images are acquired with Nikon A1Rplus using a 20x water immersion objective lens (NA=0.7) with a pixel scale of 0.243 *μm* in xy and z-step 2 *μm*. A 488 nm line of an Argon Laser was used to detect NucGreen. Note that, all 2D spheroid images were normalized to have zero mean and variance equal one to compensate for intensity variation before models training process.

### Dataset annotation

Ground truth annotation was performed in semi-automatic way using the interactive learning and segmentation toolkit (Ilastik) [53] based on shallow learning followed by expert correction (Figure 1.c). 2D frames from the beginning, middle and the end of 3D image stacks belonging to each clearing condition were selected for exhaustive representation of instances and clearing effects on image depths. Pixel wise classification mode was used then to generate primary segmentation maps. The filters used in the segmentation step were intensity, edge and textural filters with various kernel sizes ranging from 0.7 pixel to 10 pixels in order to represent the smallest and largest details of the transition between the nuclei and the background (36 filters). Then, we applied the batch processing to segment all 2D data set images. Finally, a manual correction step produced by an expert was applied to the segmentation maps. The overall amount of annotated 2D images was 57 for each clearing conditions. Datasets were then split into 47 for training, 5 for validation and 5 for testing. The total number of cell nuclei was around 10000 for each condition dataset.

3D spheroid images of various thicknesses were normalized in z-depth by considering the slice with the largest diameter as the center of spheroid. Diameter of each spheroid slice was computed from the binarized maximum intensity projection of the 3D volume on y-axis. The percentage of diameter was derived then for each slice according to the maximum diameter value. Based on this normalization method, and for a fair comparison between the protocols, we considered slices of depths ranging from 70% to 100% of maximum diameter (Figure 2A). It is to be noticed that this limited range of depth causes no loss of information since spheroid are isotropic samples. The missing part of the spheroids are similar to the first half.

### Image quality assessment based local metrics

We first assess locally the quality of spheroid images according to z-depth by using statistical metrics are the signal to noise ratio (SNR) and the contrast to noise ratio (CNR) calculated by two ways: Bhattacharyya coefficient (BC) and Fisher ratio (FR) defined as

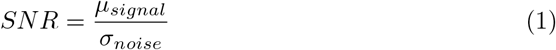

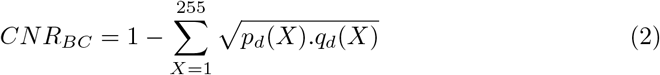

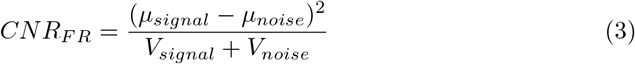

where, *μ_signal_, p_d_*(*X*) and *V_signal_* are the mean intensity of the signal, the probability of a signal intensity value *X* and the variance of the signal respectively. Also, *μ_noise_, q_d_*(*X*), *V_noise_*, and *σ_noise_* are the mean intensity of the noise, the probability of a noise intensity value *X*, the variance and the standard deviation of the noise respectively (see SI Figure 1). We computed also the normalized average intensity (average I/Imax) in each image slice to illustrate the change in intensity along the depths. All these factors were computed for three spheroids from each clearing condition.

### Deep learning segmentation methods

To assess the quality of clearing, we tested a set of state-of-the-art segmentation methods based on deep learning that were used for 2D spheroid images segmentation (Figure 1.D). They are briefly presented in this subsection.

#### U-Net

We used the reference segmentation network UNet [54] to predict 2 output classes (Cell nuclei and background). We simplified the architecture by reducing the number of feature channels. We use 5 blocks for contracting and expansive paths, each consisting of 2 convolutional layers with 4.2^*n*^ (*n* = 2, 3,4, 5, 6) filters of size 3 × 3 and ReLU activation function. For the output probability map (*ŷ*), we use a single-channel convolutional layer with sigmoid activation function (see SI Figure 6.A). The total number of trainable parameters was 2, 158, 417 optimized by using the Adam optimizer [55] and the training hyper-parameters are: *batch size* = 1, *epochs* = 33 and the learning rate *lr* = 1*e*^−3^. The final segmentation map was computed by thresholding the probability map by a threshold value *α* that was optimized on validation data set for each condition. The optimization process is described later in the same section.

#### Post processing based Dynamic Morphology (DM)

The same U-Net architecture described previously was applied to predict the probability map (*ŷ*) followed by a post processing step based on dynamic morphology [56] and watershed algorithm [57]. This combination was used in histopathology images segmentation to separate touching nuclei [58] that are considered as one object after thresholding the probability map. Briefly, the post processing step was based mainly on the hypothesis that the posterior probability at the border of the touching nuclei was systematically lower than in the putative center of the nucleus, and that the nuclei centers correspond to local maximum intensity in the image. The significant drop of the signal between nuclei center and the border was defined by morphological dynamics as the following: Let *LM* be a local maximum of the U-Net probability map output *ŷ*. *LM* represents a cell nuclei if along all paths *P* connecting *LM* with some higher maximum *LM*′, the decrease in *ŷ* is at least λ

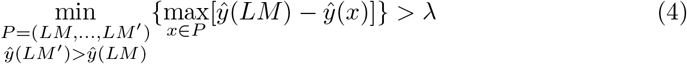

where *x* is a pixel at (*i,j*) coordinates and λ is a free parameter optimized for each clearing method (see SI Figure 6.B). The final segmentation map was then obtained by applying watershed transformation to the inverted probability map seeded from the maxima that fulfill this criterion.

#### Dist

The problem in high nuclei density 2D spheroid images is that the touching and overlapping nuclei are segmented as one object. Several works were proposed to solve this problem by predicting both the object and their contours [59, 60]. Others proposed to focus the attention of the model on the core of the nuclei by predicting an eroded version of the annotation as centers correspond to the ultimate erosion of the ground truth [61, 62]. In our work and unlike pixel-wise binary classification used previously, we follow the proposed work in [37] and we turned 2D spheroid images classification to a regression problem by predicting the distance maps 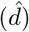 that focus on the center of nuclei (see SI Figure 6.A). Therefore, for each pixel *x* = (*i, j*) of the annotated spheroid binary image (*y*), with *y*(*x*) > 0, we assign a distance transform (*D_c_*) representing the distance to the closest background pixel *x_b_* = [*i_b_, j_b_*]. Here, we used the Chebyshev distance defined as,

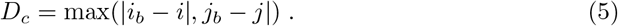

The Dist model is based on the same U-Net architecture described before. The same training hyper-parameters were used to predict distance maps. Only sigmoid output function was replaced with ReLU function at the output channel since the predicted values are higher than 1. The final binary segmentation map was then obtained by thresholding the distance output maps 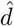. The threshold value denoted *β* was also optimized for RapiClear, Control and Glycerol clearing methods. Finally, the post processing step described before was also applied to the predicted maps to enhance the final segmentation after optimizing the parameter λ for each clearing method.

#### Stardist: Star-convex polygons

Star-convex polygons (Stardist) is one of the robust widely used algorithms for cells detection and segmentation in 2D microscopy images [36]. It consists in predicting a star-convex polygon for each cell nuclei pixel *x* = [*i, j*] by regressing the distances 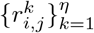 of the pixel to the boundary of the nuclei to which belong, along a set of optimized number of radial directions *η* with equidistant angles. Also, separately, the algorithm predicts probability map (*p_i,j_*) for each pixel *x* as the normalized Euclidean distance to the nearest background pixel *x_b_* = [*i_b_, j_b_*] (see SI Figure 6.A). Given such polygon candidates with their associated nuclei probabilities, a non-maximum suppression (NMS) was performed to reach the final set of polygons, each representing a cell nuclei. Stardist was mainly based on Unet architecture with 3 blocks for contracting and expansive paths, each consisting of 2 convolutional layers with 32.2^*n*^ (*n* = 0,1, 2) filters of size 3 × 3, and an additional layer of 128 3 × 3 filters added after the final Unet feature layer to avoid that the subsequent two output layers have to fight over features. The activation functions between layers are ReLU and the total number of trainable parameters was approximately 1.4 million parameters. The output layers of the architecture consist of a single-channel convolutional layer with sigmoid activation for the nuclei probability output and the polygon distance output layer has as many channels as there are radial directions *η* and do not use an additional activation function. We used Stardist in our study to segment cell nuclei inside 2D spheroid images. For the three conditions datasets, the training stage was performed with the following primary hyper parameters: *batch size* = 1, *epochs* = 400 and the learning rate *lr* = 0.3*e*^−3^. After that, the best training model that minimized the loss functions (Equations 6, 7) was selected for each condition. So, based on this criterion the hyper parameters for the selected models of RapiClear, Control and Glycerol datasets were (*epochs* = 317, *lr* = 0.75*e*^−4^), (*epochs* = 282, *lr* = 1.5*e*^−4^) and (*epochs* = 380, *lr* = 0.75*e*^−4^) respectively. As in the previously described segmentation methods, theparameters: number of rays *η*, threshold *α* of the nuclei probability map and threshold of the non-maximum suppression *τ*, were optimized for each data set.

### Training loss functions and segmentation evaluation metrics

#### Loss function

The purpose of loss function in a deep learning training stage model is to quantify the difference between predictions and ground truths for steering the training of the network. In our work, we used two commonly used loss functions the binary cross entropy (*BCE*) and the mean squared error (*MSE*) for classification and regression problem respectively. The *BCE* for binary classification (nuclei and background) was defined as

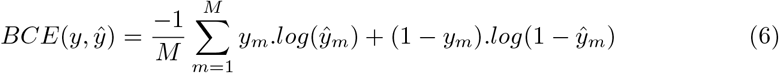

and *MSE* for distance map prediction was defined as,

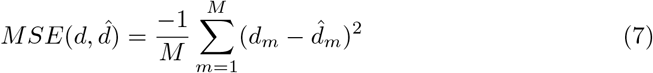

where *M* is the 1*D* output map size, *ŷ_m_* and 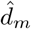 are the m-th scalar values in the output predicted maps, *y_m_* and *d_m_* the corresponding target values of probability maps and distance maps respectively.

#### Evaluation metrics

To evaluate the performance of spheroid images segmentation and to be able to quantitatively compare between the transferability of segmentation models trained on clarification method to other clearing protocols, we used the pixel wise-metric F1-score [63] for segmentation evaluation and also the Aggregated Jaccard Index (AJI) [61] as an object-wise metric for touching nuclei splitting evaluation. The F1 measure is defined as the harmonic mean between recall and precision at the pixel level and it was computed as

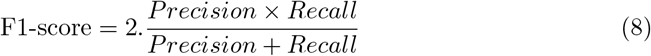

with the 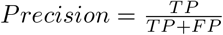 and 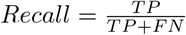, where *TP*, *FP* and *FN* are the true positive, false positive and false negative respectively.

The AJI is an extension of the global Jaccard index, where every ground truth nucleus is first matched to one detected nucleus by maximizing the Jaccard index. The AJI corresponds then to the ratio of the sums of the cardinals of intersection and union of these matched components respectively. In addition, all detected components that do not matched were added to the denominator. More formally [37], AJI can be defined as,

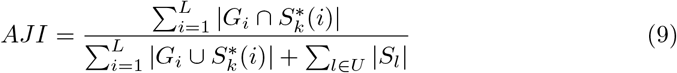

where *G_i_* is a nucleus ground truth of *L* nuclei in an image, *S* is all the detected nuclei, 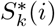 is the segmented nucleus associated with *G_i_* that maximizes the Jaccard index, i.e. 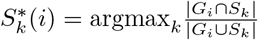 and *U* is the set of indices of detected nuclei that have not been assigned to any ground truth.

### Parameter Optimization

One important step in supervised learning with deep learning is the hyperparameters fine tuning. This optimization step was applied to validation data set to maximize the segmentation metrics such as F1-score, AJI or Jaccard index for Stardist segmentation method. In our work, threshold of the probability maps *α*, threshold of predicted distance maps *β*, the h-minima value λ of the dynamic morpholoy reconstruction post processing step, the numbers of rays of the star-convex polygons *η* and threshold *τ* of the non-maxmimum suppression (nms) were optimized empirically by varying each parameter between a range of values (see SI Figure 7). Then, the value that maximized a segmentation metric were selected for each segmentation method depending on the RapiClear, Control and Glycerol segmentation models as shown in Table 2.

**Table 2.** Parameters optimization step. Various thresholds were optimized for each used segmentation method between a range of tested values. The value of threshold that maximized a metric was selected.

| Parameters | Segmentation methods | Ranges | Metrics | Segmentation models |  |  |
| --- | --- | --- | --- | --- | --- | --- |
|  |  |  |  | *Control* | *RapiClear* | *Glycerol* |
| $\alpha$ | Unet | $\alpha \in [0 \ 1]$ | F1-score | 0.5 | 0.6 | 0.4 |
| | Stardist | $\alpha \in [0.3 \ 0.7]$ | Jaccard index | 0.6 | 0.4 | 0.5 |
| $\beta$ | Dist | $\beta \in [0 \ 2]$ | F1-score | 1.1 | 1.2 | 0.6 |
| $\lambda$ | Unet + DM | $\lambda \in [0 \ 20]$ | AJI | 8 | 7 | 2 |
|  | Dist + DM |  |  | 2 | 1 | 1 |
| $\eta$ | Stardist | $\eta \in 2^i, \ i \in [2 \ 8]$ | F1-score | 64 | | |
| $\tau$ | | $\tau \in [0.3 \ 0.5]$ | Jaccard index | 0.4 | 0.3 | 0.4 |

### Transfer learning

We used transfer learning [64] methodology in our study to transfer knowledge gained from training segmentation model on a dataset from a clarification condition and apply the pre-trained model to segment images clarified from other conditions. This was done by brute transfer of the weights in the neural network without fine tuning.

## Supporting information

**S1 Fig. Supplementary Figures 1,2,3,4,5 and 6**

**S1 Datasets and Videos. Datasets availability and 3D spheroids reconstruction videos.** Annotated data sets used during the current study are available and supplementary videos of 3D reconstructed spheroids for Control (non cleared), Rapiclear and Glycerol cleared samples are available in the following repository: https://uabox.univ-angers.fr/index.php/s/6myuGGs0JO94M8D.

## Acknowledgments

Ali Ahmad gratefully acknowledges project EU H2020 FET Open, PROCHIP, Chromatin organization PROfiling with high-throughput super-resolution microscopy on a CHIP, grant agreement no. 801336 (https://pro-chip.eu/) for funding his PhD. The authors also thank the GdR ImaBio for financial support to the organisation of a deep learning hackathon which contributed to the initiation of this study.

## Author contributions statement

A.A. and D.R. conceived of and designed the experiments. S.G., G.R. and C.R prepared and acquired the spheroid samples. A.A. annotated the data used in the experiments. A.A conceived and developed image quality assessment and image segmentation based on deep learning algorithms. A.A., D.R. G.R. and C.R analysed the data and the results. All authors contributed to the redaction of the manuscript and approved the final version. D.R. and C.F. supervised the PROCHIP project.

---

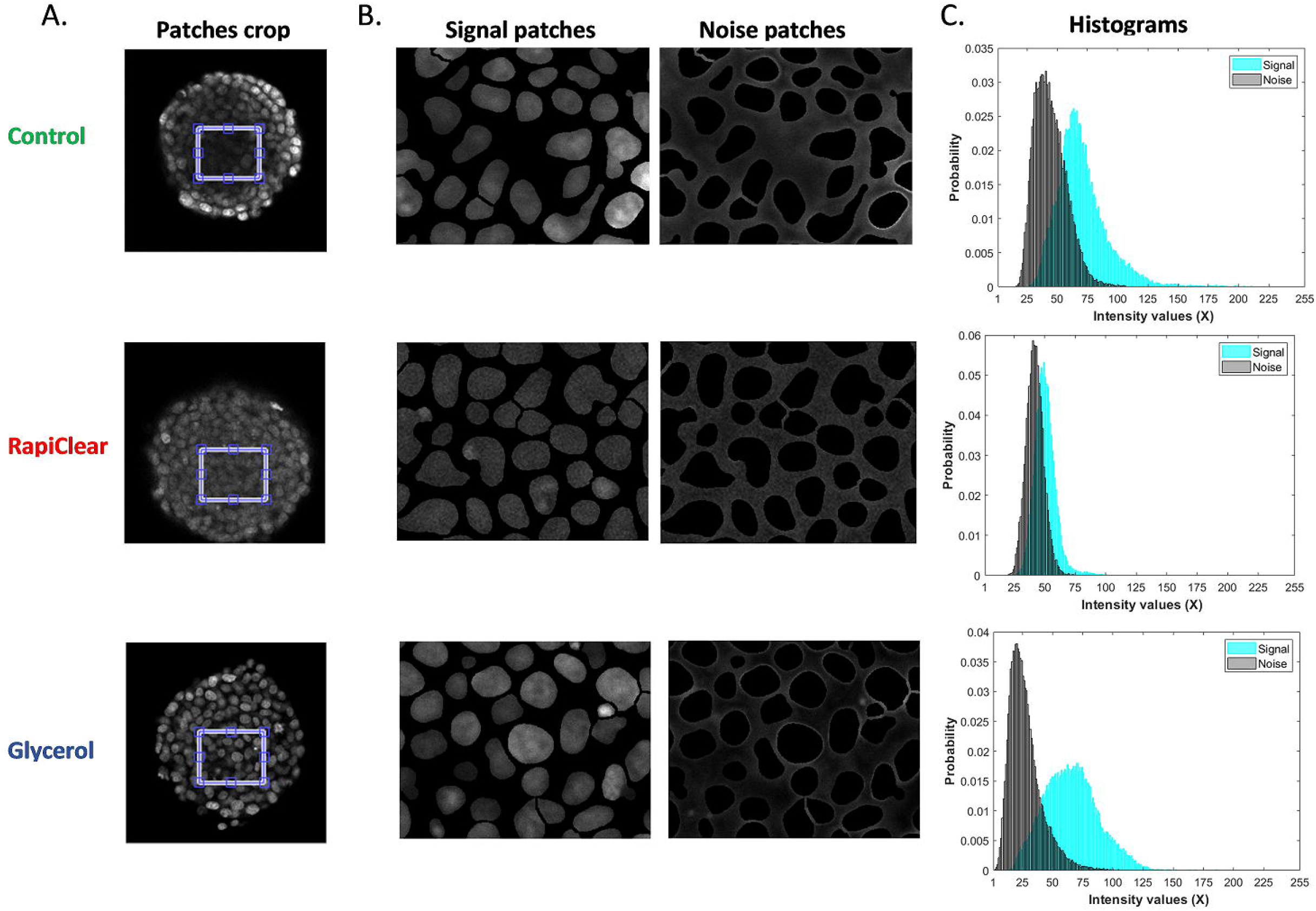

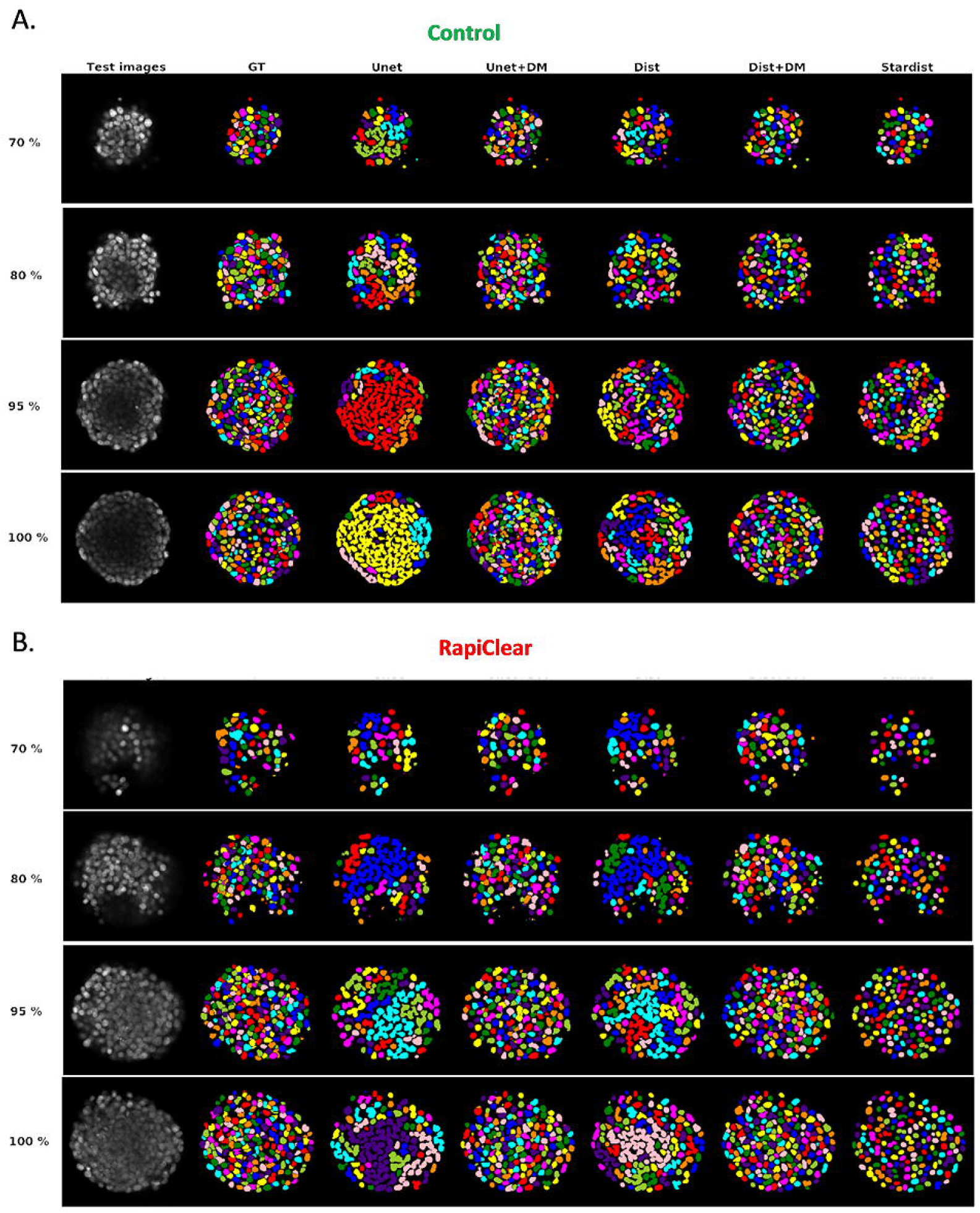

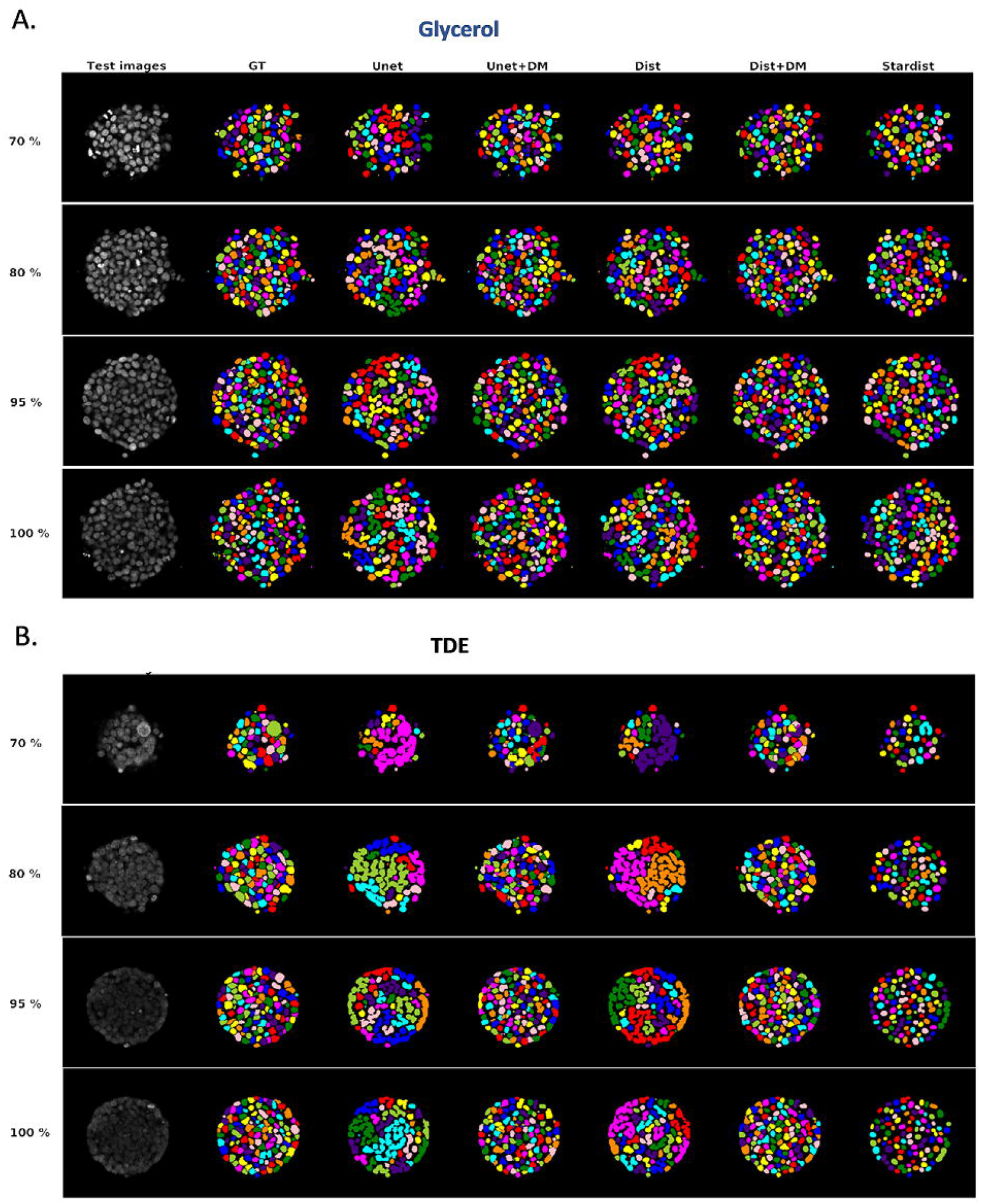

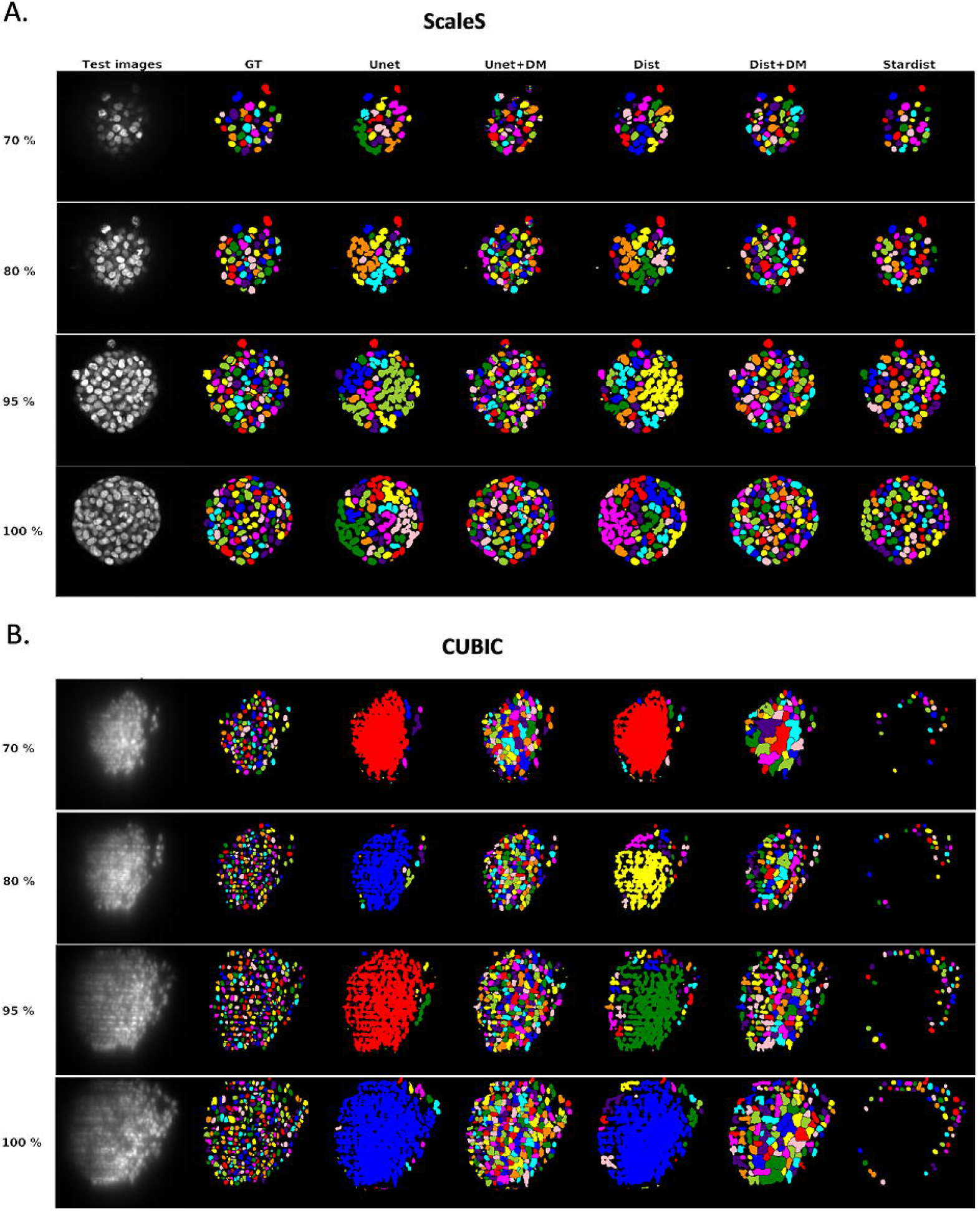

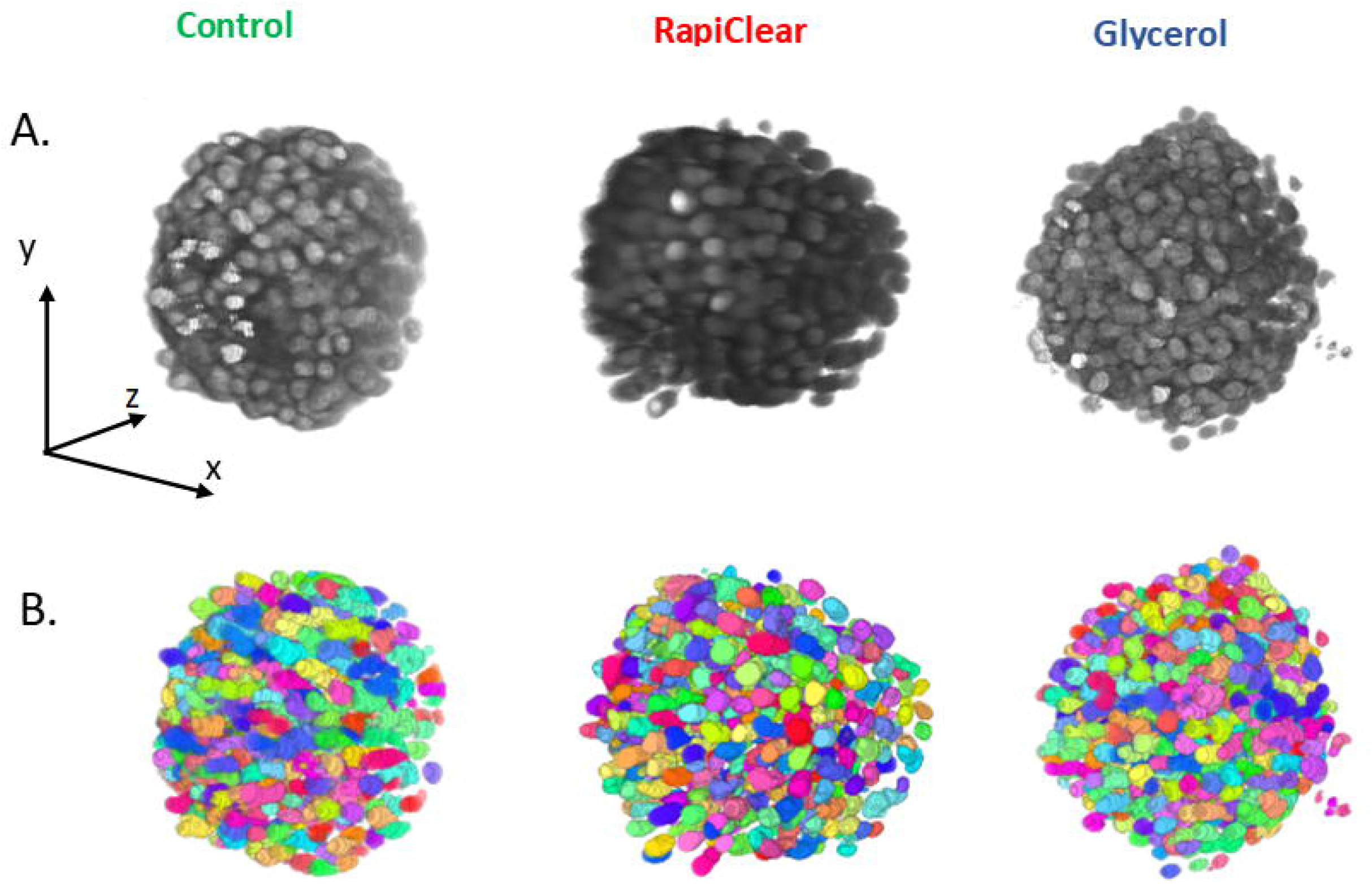

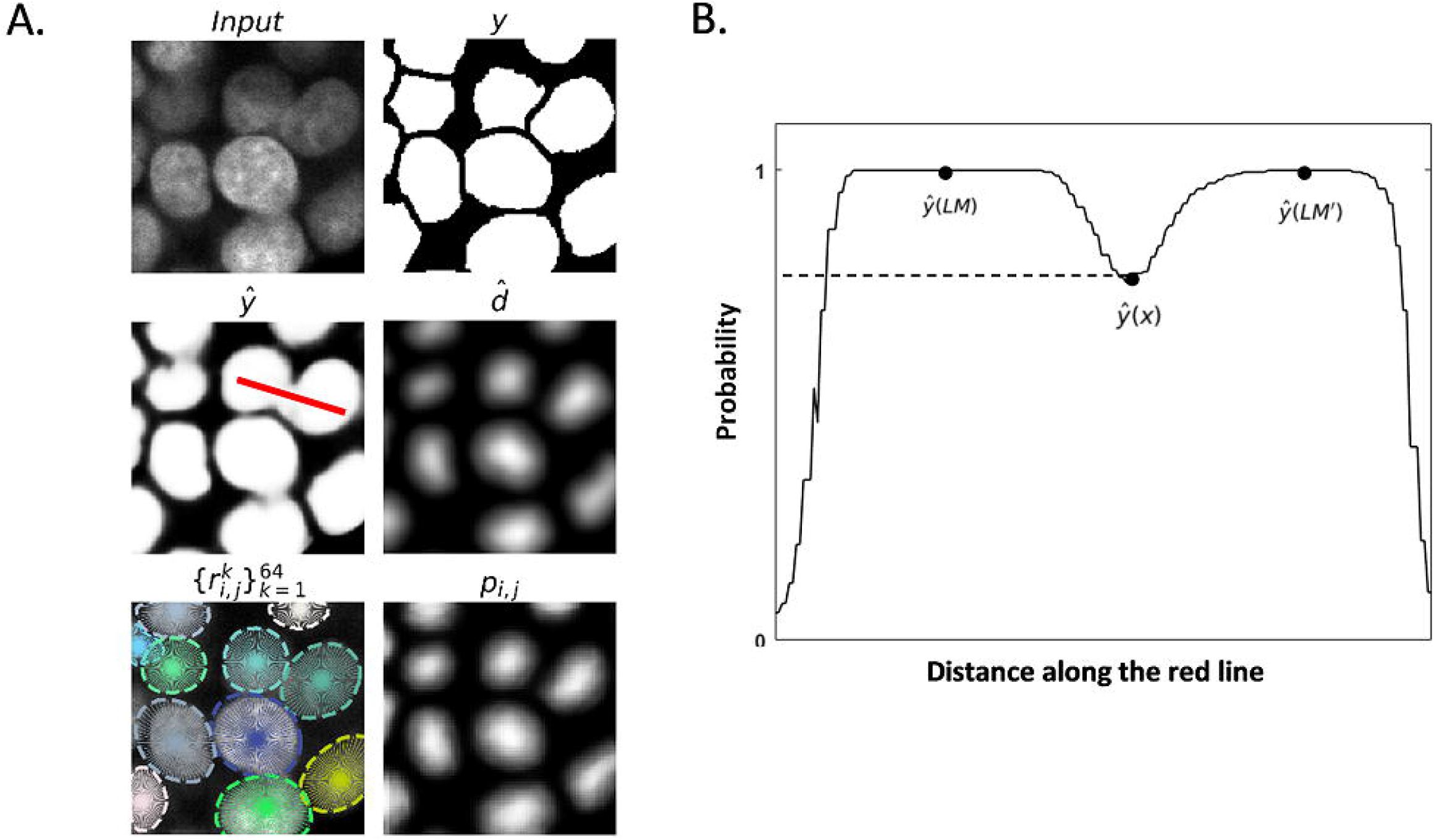

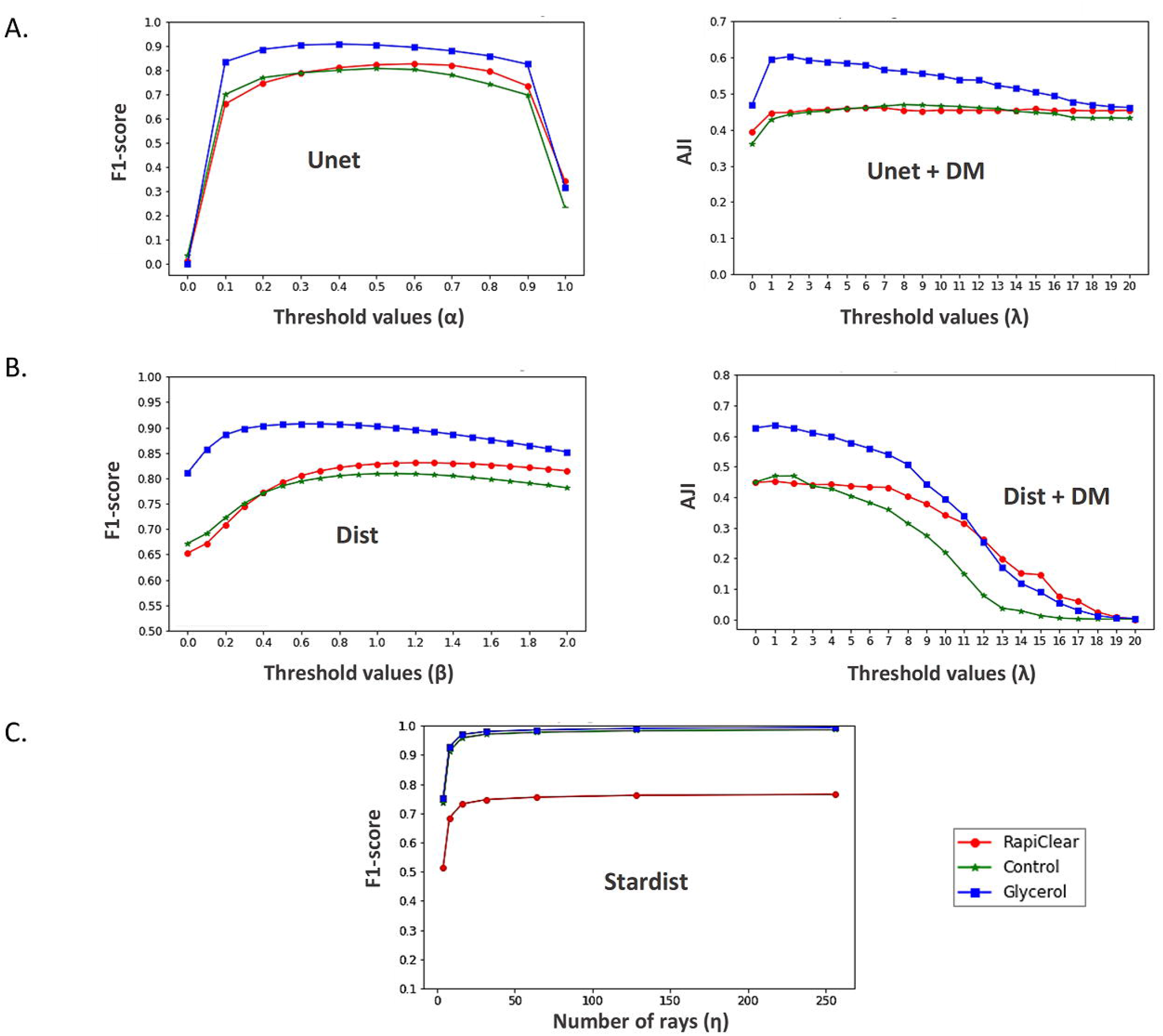

## Notes

https://uabox.univ-angers.fr/index.php/s/6myuGGs0JO94M8D

### Competing Interest Statement

The authors have declared no competing interest.

