## Supplementary figures 1,2,3,4,5,6 and 7 for "Clearing spheroids for 3D fluorescent microscopy: combining safe and soft chemicals with deep convolutional neural network"

### Supporting information

#### S1 Figure 1

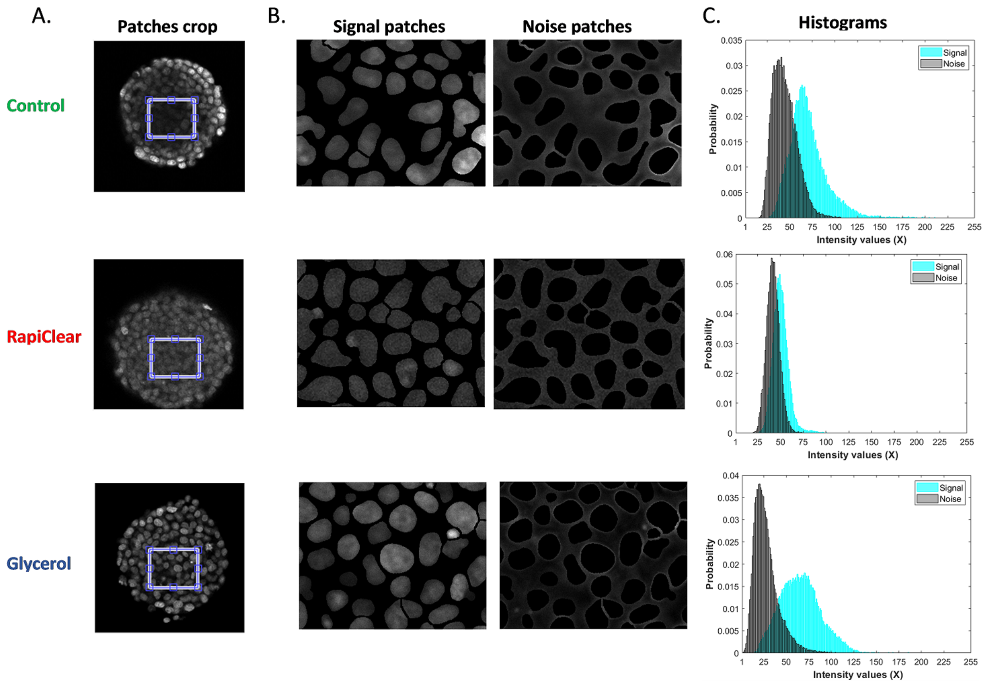


**Fig 1. Definition of patches for the computation of local metrics. (A)** Patches are cropped from the center of spheroid in each slice along z-depth for Control, RapiClear and Glycerol datasets. **(B)** Each patch is split into nuclei taken as signal and noise as the extracellular background. They are used to compute signal to noise ratio (SNR) and contrast to noise by Fisher ratio ($CNR_{FR}$). **(C)** Histograms illustrate the probability values of (X) intensity, where $X \in [1255]$ of nuclei signal and noise. These distributions are used to compute contrast to noise ratio by Battacharayya coefficient ($CNR_{BC}$).

#### S1 Figure 2

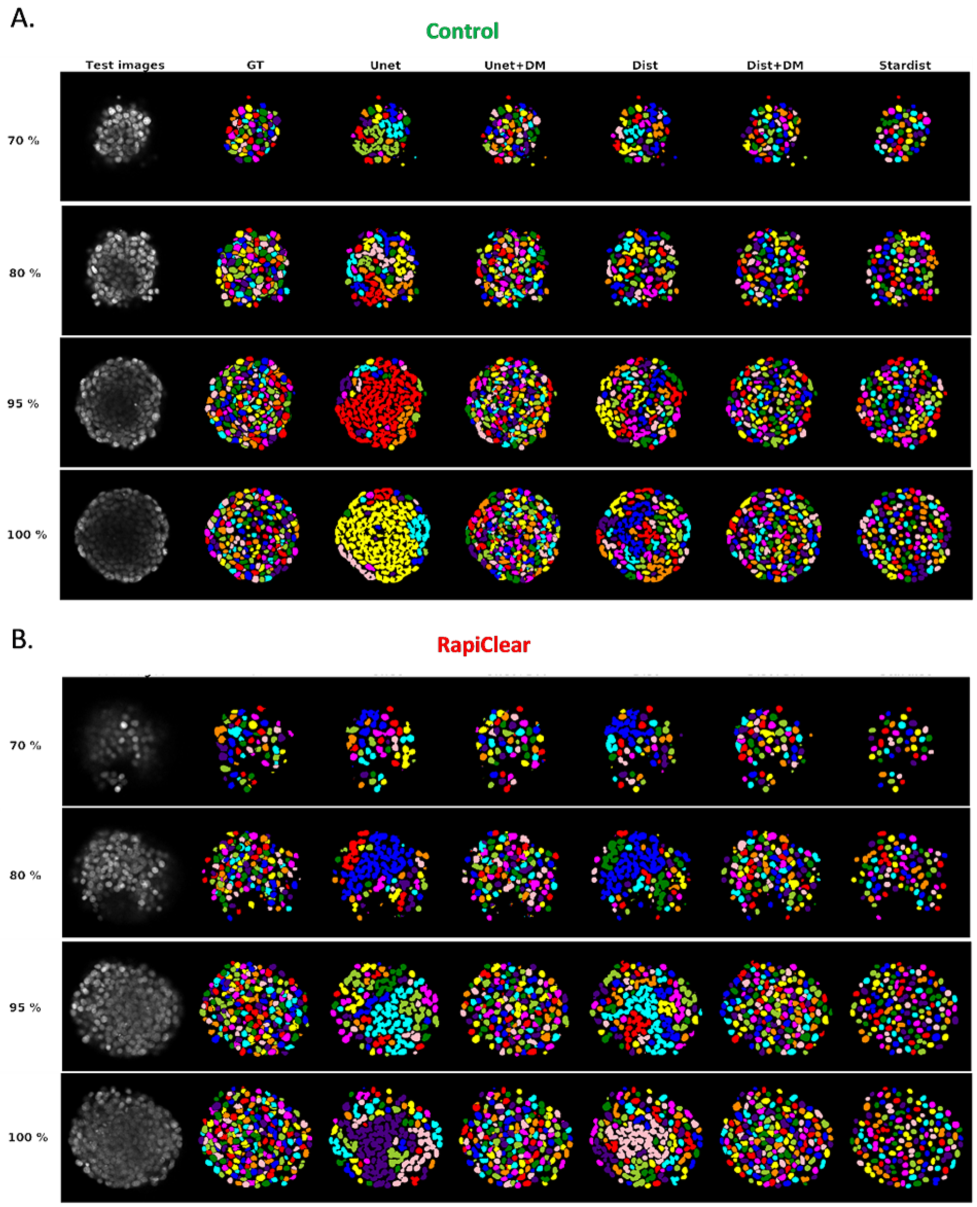


**Fig 2. Final segmentation maps for control and RapiClear images.** Illustration of the final segmentation results of all used segmentation methods for the slices at 70, 80, 95 and 100% of maximum diameter (z-depth) **(A)** for Control and **(B)** RapiClear test mages.

#### S1 Figure 3

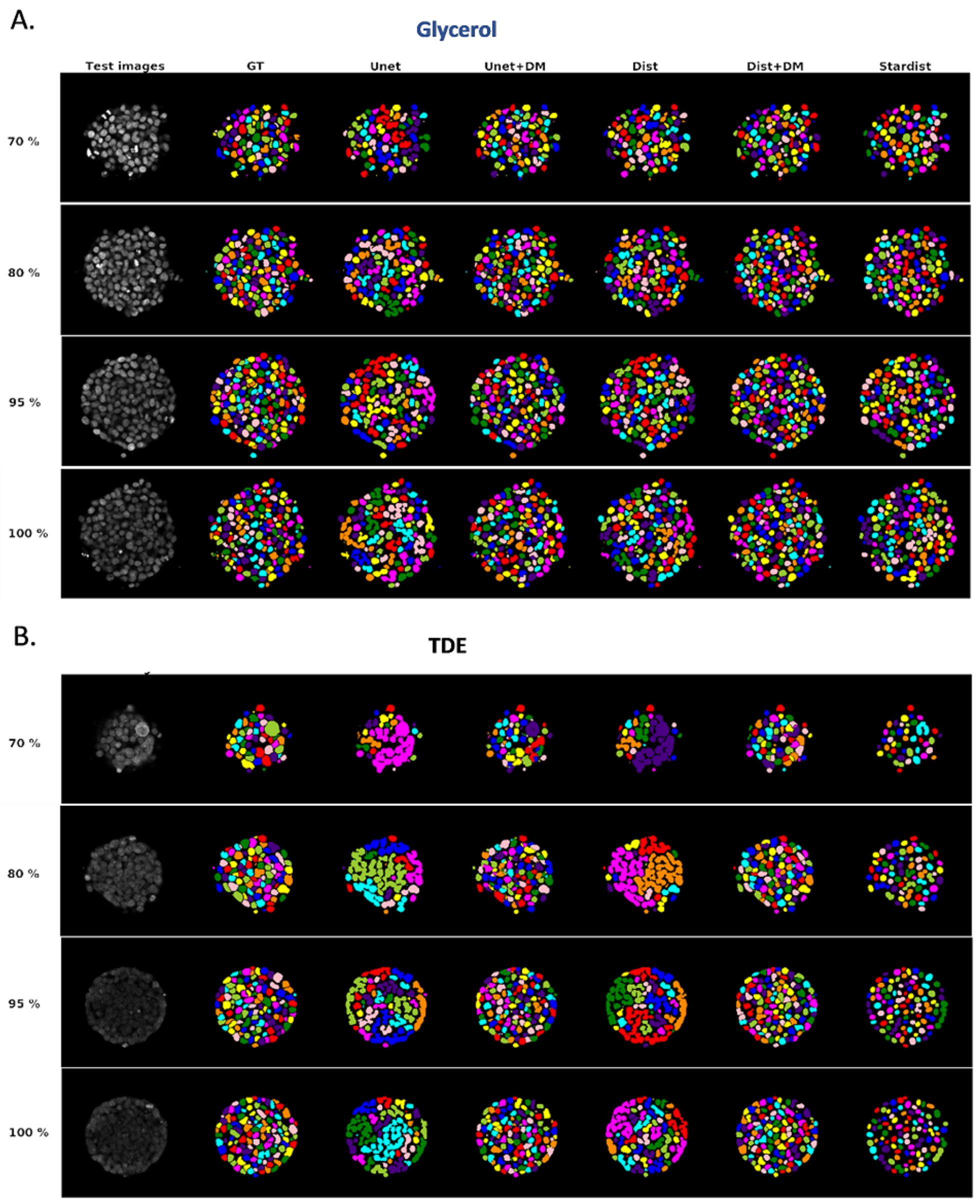


**Fig 3. Final segmentation maps for Glycerol and TDE images.** Illustration of the final segmentation results of all used segmentation methods for the slices at 70, 80, 95 and 100% of maximum diameter (z-depth) **(A)** for Glycerol test images and **(B)** TDE test images segmented using RapiClear model during data transferability test.

#### S1 Figure 4

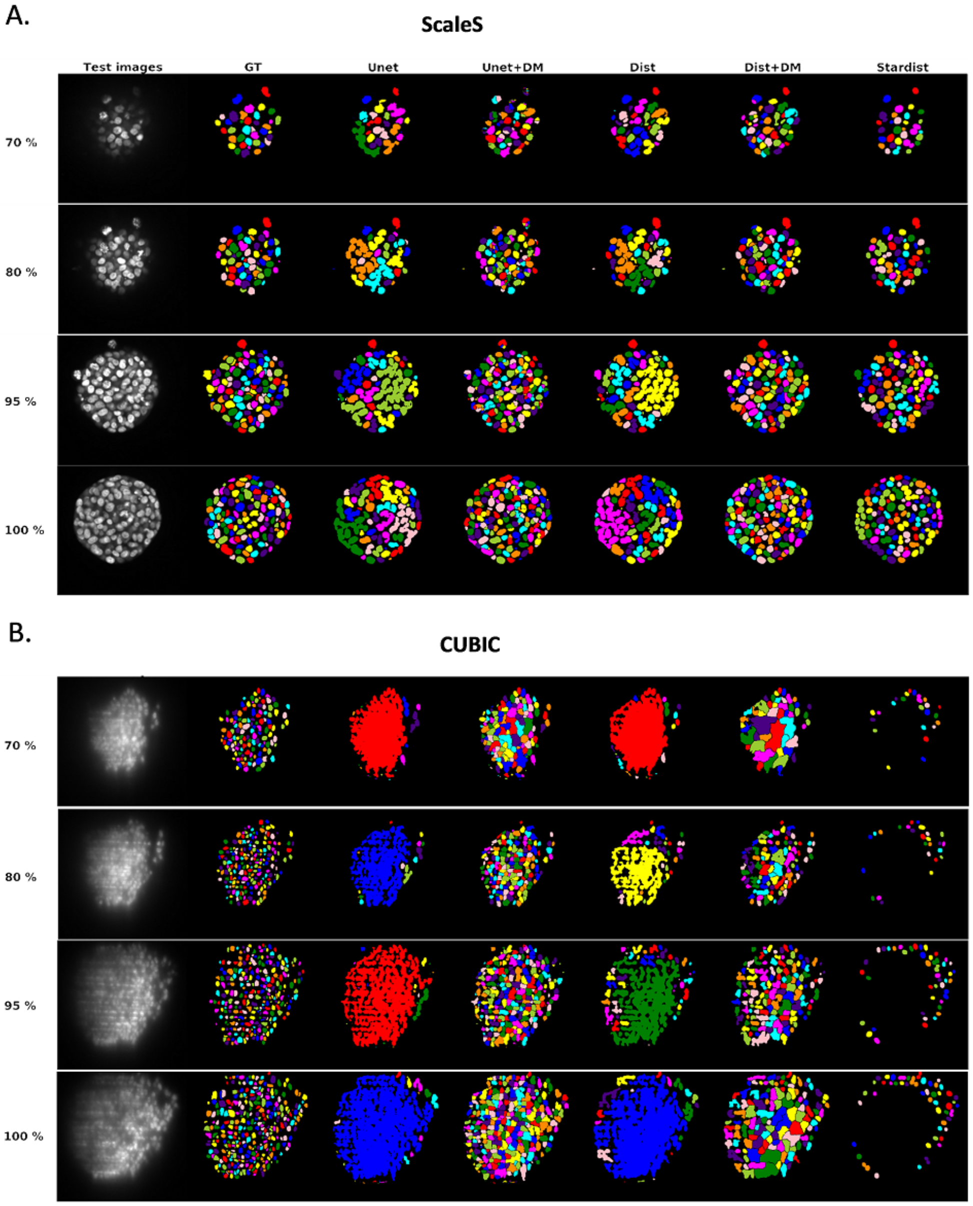


**Fig 4. Final segmentation maps for ScalesS and CUBIC images.** Illustration of the final segmentation results of all used segmentation methods for the slices at 70, 80, 95 and 100% of maximum diameter (z-depth) **(A)** for ScaleS test images segmented using Glycerol model and **(B)** CUBIC test images segmented using Control model during data transferability test.

#### S1 Figure 5

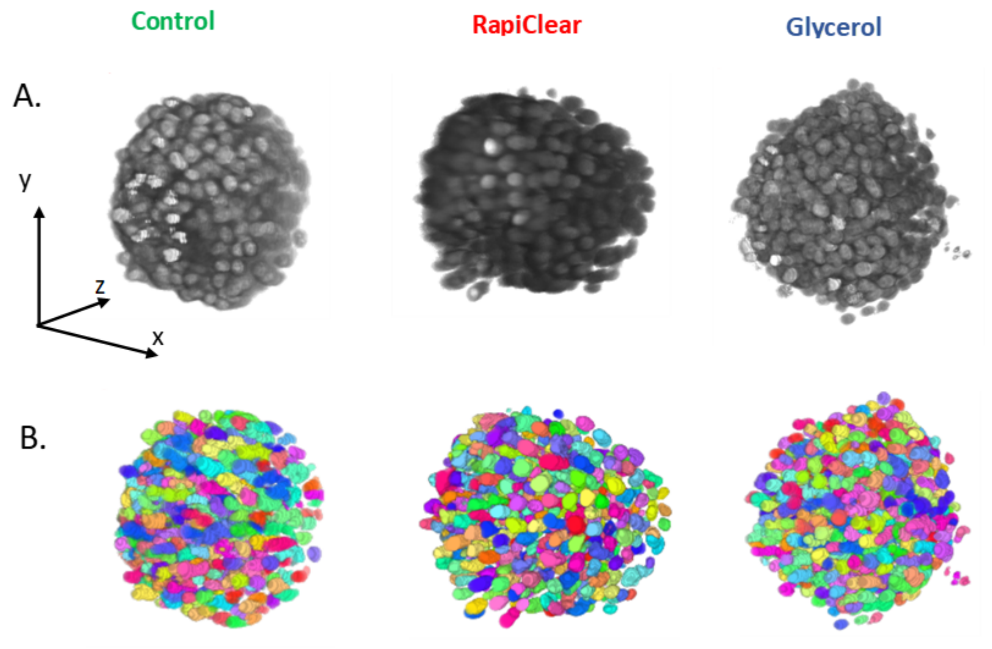


**Fig 5. Spheroids 3D reconstruction. (A)** Control, RapiClear and Glycerol spheroid samples used for 2D segmentation (figure 2 in the main manuscript) with a Glycerol pre-trained Dist model. **(B)** Illustration of the 3D reconstruction of each spheroid after 2D segmentation.

#### S1 Figure 6

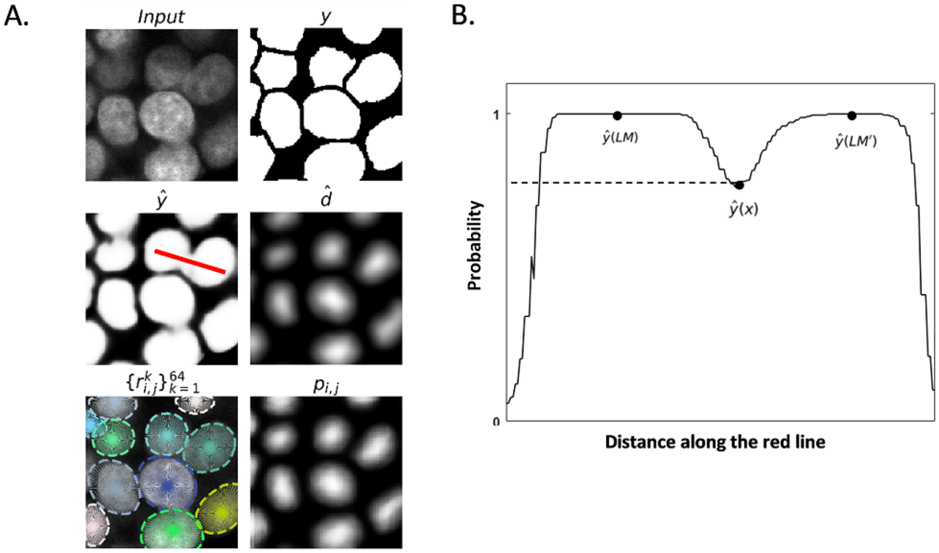


**Fig 6. Deep learning segmentation method outputs. (A)** Illustration of a cropped patch from 2D spheroid image and its ground truth binary mask ($y$). When segmented with Unet the output is the inversed probability map $\hat{y}$. $\hat{d}$ is the predicted distance map at the output of Dist. $\{r_{i,j}\}_{k=1}^{64}$ and $P_{i,j}$ are the predicted distances of a pixel $x = (i, j)$ to the boundary of nuclei along a set of radial directions (empirically found optimal at $\eta = 64$) and the object probability map at the output of Stardist respectively. **(B)** Illustration of the principle of post processing-based dynamic morphology reconstruction.

#### S1 Figure 7

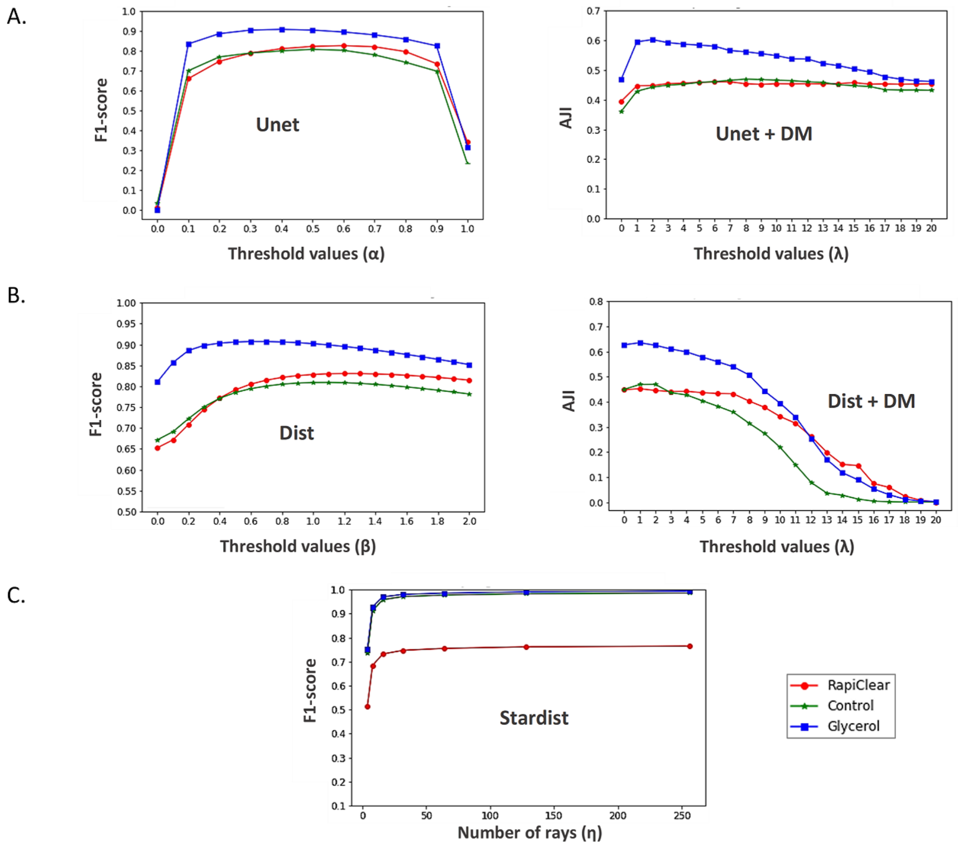


**Fig 7. Hyperparameters optimization based on validation dataset for Control, RapiClear and Glycerol segmentation models.** Plots shows the evolution of $F1 - score$ and $AJI$ depending on the values of **(A)** probability map threshold $\alpha$ and h-minima parameter $\lambda$ for Unet and Unet+DM respectively, **(B)** the predicted distance map threshold $\beta$ and h-minima value $\lambda$ for Dist and Dist+DM and **(C)** the variation of $F1 - score$ metric function of the number rays $\eta$ required to reconstruct the polygons.
